# Transcriptome-wide spatial RNA profiling maps the cellular architecture of the developing human neocortex

**DOI:** 10.1101/2021.03.20.436265

**Authors:** Kenny Roberts, Alexander Aivazidis, Vitalii Kleshchevnikov, Tong Li, Robin Fropf, Michael Rhodes, Joseph M. Beechem, Martin Hemberg, Omer Ali Bayraktar

## Abstract

Spatial genomic technologies can map gene expression in tissues, but provide limited potential for transcriptome-wide discovery approaches and application to fixed tissue samples. Here, we introduce the GeoMX Whole Transcriptome Atlas (WTA), a new technology for transcriptome-wide spatial profiling of tissues with cellular resolution. WTA significantly expands the Digital Spatial Profiling approach to enable *in situ* hybridisation against 18,190 genes at high-throughput using a sequencing readout. We applied WTA to generate the first spatial transcriptomic map of the fetal human neocortex, validating transcriptome-wide spatial profiling on formalin-fixed tissue material and demonstrating the spatial enrichment of autism gene expression in deep cortical layers. To demonstrate the value of WTA for cell atlasing, we integrated single-cell RNA-sequencing (scRNA-seq) and WTA data to spatially map dozens of neural cell types and showed that WTA can be used to directly measure cell type specific transcriptomes *in situ*. Moreover, we developed computational tools for background correction of WTA data and accurate integration with scRNA-seq. Our results present WTA as a versatile transcriptome-wide discovery tool for cell atlasing and fixed tissue spatial transcriptomics.

## Introduction

Spatial genomic technologies enable the analysis of the transcriptome and other molecular information *in situ*^1^. While spatial transcriptomic approaches are generally used to validate observations from single-cell RNA-sequencing (scRNA-seq), they can potentially serve as powerful discovery tools to explore human tissue architecture and pathology. To discover diverse cell states and spatiotemporal gene expression programmes regulating tissue development and function, it is essential that spatial technologies assay the whole transcriptome. Similarly, transcriptome-scale detection is required to characterise novel pathological processes in patient-derived tissue samples in situ. However, many existing spatial transcriptomic technologies are limited in assaying the whole transcriptome or poorly compatible with fixed clinical samples.

Amongst spatial transcriptomic approaches, imaging methods based on cyclic RNA hybridisation (e.g. MER-FISH, ISS) can quantify mRNA transcripts at single-cell resolution in tissues, but are often limited to a few hundred pre-selected genes^2,3^. While thousands of transcripts can be resolved by performing high numbers of cycles^4^, the increased imaging time limits assay throughput to small tissue areas. Alternatively, spatially resolved RNA-seq methods that utilise array-based mRNA capture from tissues (e.g. 10x Visium, Slide-sequencing) can provide unbiased transcriptome-wide information at high throughput^5–7^. However, spatial RNA-seq methods sacrifice cellular resolution as the arrays capture multiple cell types or anatomical structures at each spatial position. Moreover, RNA-seq methods are poorly applicable to formalin-fixed paraffin-embedded (FFPE) material that constitutes the majority of clinical tissue samples.

The GeoMx Digital Spatial Profiling (DSP) method combines the strengths of imaging and sequencing approaches for spatial transcriptomic analysis^8^. DSP uses RNA hybridisation probes or antibodies labelled with photocleavable next generation sequencing (NGS) indexing tags. These indexing tags are captured from specific cell types or regions on tissue sections using microscopy-guided photocleavage, and quantified using NGS. Hence, DSP achieves high-throughput quantification of mRNA targets with cell type resolution. Importantly, DSP works optimally on FFPE tissue material owing to probe hybridisation based mRNA detection. However, to date, DSP has been used to detect only up to 1,811 genes in a multiplexed manner *in situ*^9–11^ *which is insufficient for unbiased characterization of complex tissues*.

The fetal human neocortex shows a remarkable degree of spatial cellular organisation that underlies cortical development and function. Across the fetal cortex, diverse progenitor and differentiating cell types are organised into distinct cellular compartments along the cortical depth^12^. Multiple types of radial glial stem cells and intermediate progenitors reside in the deep germinal zones. These neural progenitors generate excitatory neuron and glial subtypes that migrate towards the superficial cortical plate and form the six layers of the adult neocortex. Despite extensive histological^13,14^ and scRNA-seq^15,16^ characterisation of fetal cortical cell types, we lack a comprehensive spatially resolved analysis of neocortical cellular architecture.

Here, we introduce the GeoMX Whole Transcriptome Atlas (WTA), a new technology for transcriptome-wide spatial profiling of fixed tissue samples. WTA significantly expands the DSP approach to enable *in situ* hybridisation against 18,190 genes at high-throughput using a sequencing readout. We apply WTA to generate the first spatial transcriptomic map of the fetal human neocortex, validating transcriptome-wide *in situ* profiling of FFPE tissue material and demonstrating the spatial enrichment of autism gene expression in deep cortical layers. To demonstrate the value of WTA for cell atlasing, we integrate scRNA-seq and WTA data to spatially map dozens of neural cell types. Moreover, we present new computational tools for background correction of WTA data and accurate integration with scRNA-seq. Finally, we show that WTA can directly measure cell type specific transcriptomes *in situ*.

## Results

### WTA profiling of the fetal human neocortex

The WTA technology combines transcriptome-wide *in situ* hybridisation with a NGS readout to quantify spatial gene expression in tissue. To profile specific tissue features (e.g. anatomical regions) or cell types *in situ*, WTA is integrated into the DSP workflow^8^ (Fig. 1a). First, FFPE tissue sections are fluorescently labelled with ‘visualisation markers’ via RNA fluorescent in situ hybridisation (FISH) or antibody staining select of tissue features or cell types. Second, WTA *in situ* hybridisation probes targeting 18,318 human transcripts (Supp. Table 1) are applied to tissues. Each probe is tagged with a photocleavable unique molecular identifier (UMI) for NGS. The WTA panel also includes 138 negative control probes against ERCC sequences to estimate non-specific probe binding. Third, labelled tissue slides are processed on the DSP instrument, whereby users are guided by the visualisation markers to photocleave and collect WTA probe tags from selected regions of interest (ROIs). These ROIs are flexibly defined to target specific tissue features or cell types via manual or automated image segmentation. Finally, the WTA probe UMI counts from each ROI are obtained using NGS to provide spatially resolved transcriptome data.

**Figure 1:**
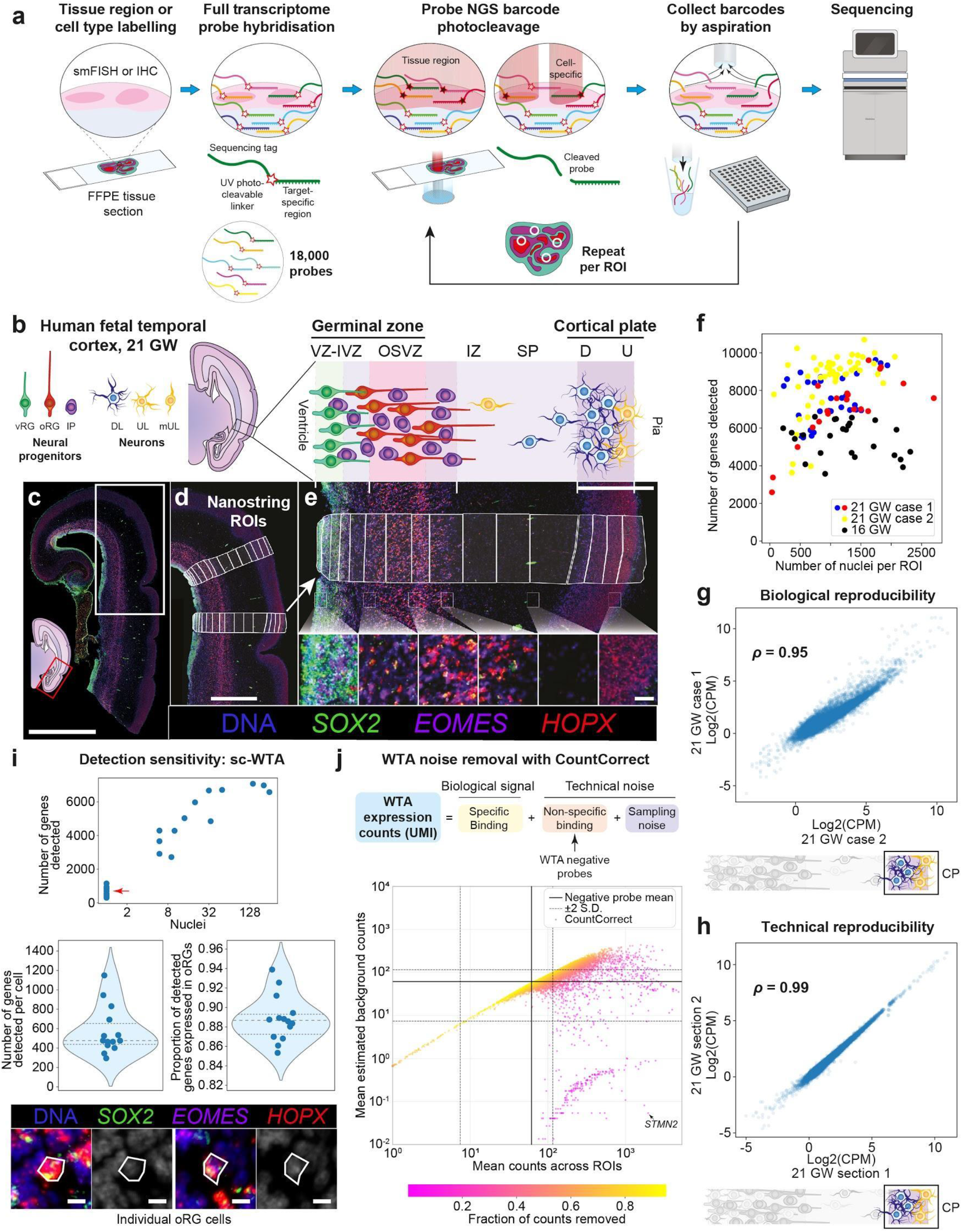
WTA in the fetal human brain. **a**, NanoString WTA experimental workflow. **b**, Diagrams illustrating the cellular architecture of the developing temporal neocortex. **c-e**, Experimental design shown for one 21 GW case. **c**, Fetal brain section stained for canonical cell type markers with RNAscope smFISH. White box shows the area enlarged in **d. d**, WTA bulk ROIs spanning the cellular differentiation and migration trajectory from the ventricle to the cortical plate. **e**, Close-up views of one strip of ROIs across the cortical depth. **f**, Gene expression detection sensitivity in bulk ROIs. A limit of detection (LoD) threshold, defined for each ROI as the mean plus 2 standard deviations of the negative probe counts, was used. The two ROIs with lowest gene counts are from the subplate, which is more sparsely populated with cells than other tissue areas. **g**, Biological reproducibility, comparing transcriptome-wide expression between the most superficial cortical plate ROI in two 21 GW cases. **h**, Technical reproducibility, comparing transcriptome-wide expression between the most superficial cortical plate ROI in two adjacent sections of the same 21 GW case. **i**, Low input and sc-WTA. oRG cells marked by SOX2 and HOPX RNAscope smFISH, were targeted by ROIs of differing sizes in the OSVZ in order to assess detection sensitivity near and at single-cell level. (Top) across the range of 1-130 cells, the number of genes detected is correlated with the number of cells targeted. Even in single oRG cells (red arrow) (example images at bottom), 300-1200 genes were detected per cell: (Middle, left) violin plot showing number of genes detected, using the same LoD as in **f**. (Right) violin plot showing proportion of genes detected that were expressed in a scRNA-seq reference oRG transcriptome. Horizontal lines show quartiles. **j**, CountCorrect noise removal model, which separates WTA counts into biological signal and technical noise. Shown is the relationship between total counts for a gene and counts assigned to background, compared to the average negative probe counts. Scale bars: 5 mm (**c**), 2 mm (**d**), 1 mm (**e**, top panel), 50 μm (**e**, bottom panels), 10 μm (**i**).

To validate transcriptome-wide spatial profiling by WTA, we profiled the cellular compartments of the developing human neocortex spanning the differentiation trajectory of cortical cells (Fig. 1b). We assayed the temporal neocortex at 16 and 21 gestational weeks (GW), distinguishing cellular compartments on FFPE tissue sections based on anatomical landmarks and cell markers defined by smFISH: *SOX2* labels neural progenitors with an enrichment in the ventricular zone (VZ), *HOPX* marks outer radial glia (oRGs) in the outer sub-ventricular zone (OSVZ), and *EOMES* labels intermediate progenitors (IPs) across both areas^17^. We profiled segmental bulk ROIs along the cortical depth, sampling across the germinal zones and the cortical plate (Fig. 1c-e, Supp. Fig. 1).

Our WTA ROIs ranged between 0.012 and 0.39 mm^2^ in area and sampled between 35 and 2,700 cells (median 1,070) as quantified by nuclear segmentation of DSP images (Supp. Fig. 2a). We detected the expression of 5,000 to 10,000 genes in the majority of ROIs across our samples (Fig. 1f) using a conservative detection threshold based on negative probe counts (Methods). The total number of genes detected with WTA was comparable to that previously detected with scRNA-seq in the mid-gestational cortex^16^ (Supp. Fig. 2b). The WTA probe counts were highly reproducible between different donors (Fig. 1g) and consecutive tissue sections (Fig. 1h). These results show that WTA achieves transcriptome wide RNA detection *in situ* with high reproducibility.

**Figure 2:**
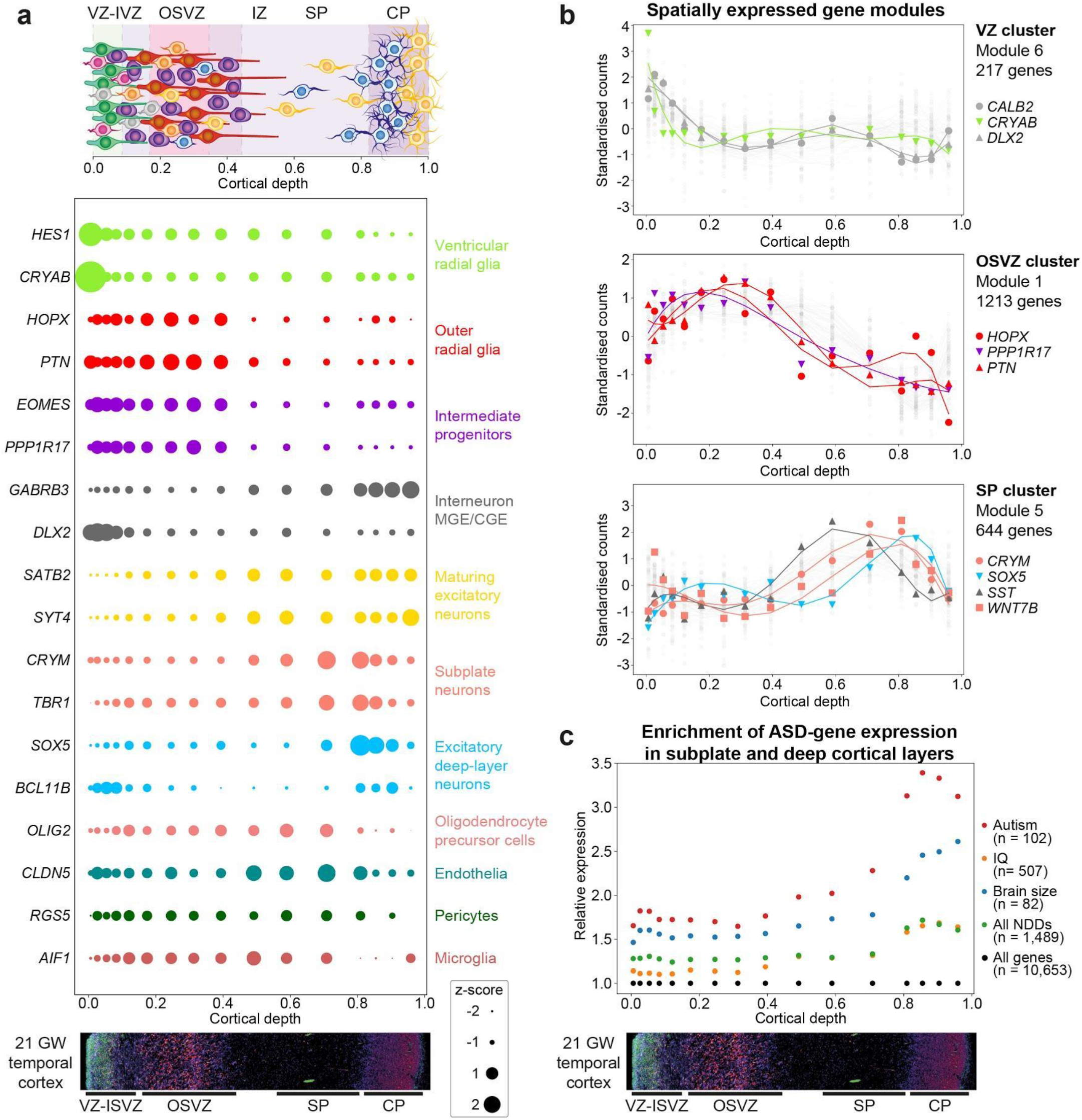
WTA accurately maps spatial cell type markers and autism-related gene expression in the fetal cortex. **a**, z-scored expression of canonical cell type markers in bulk ROIs across the depth of the temporal cortex, shown for one technical replicate of 21 GW case 1. **b**, Modules of spatially varying genes identified with SpatialDE using both technical replicates of 21 GW case 1. Three out of 12 modules are shown here. Highlighted in each are cell type markers showing the alignment of the expression profiles of these modules with known cortical compartments and resident cell types, namely: vRGs and interneurons in VZ; oRGs in OSVZ; and subplate and deep cortical neurons in SP-CP. Shown in light grey are 70 other genes in each module. **c**, Enrichment of ASD gene expression in subplate and deep cortical layers, using both technical replicates of 21 GW case 1. The average expression of four gene sets -ASD^22^, brain size^23^, IQ^24^, and all neurodevelopmental disorders^25^ -was divided by the average expression of all genes in each ROI to obtain a relative expression score and spatial enrichment patterns for each. LoD was calculated by the mean plus 2 standard deviations of the negative probe counts (10,653 genes total).

To determine the sensitivity of WTA for single-cell detection, we profiled different quantities of oRG cells from the OSVZ (Fig. 1i). In manually selected single oRG cells, we detected the expression of 574 genes per cell on average, these genes overlapped highly with known markers obtained from scRNA-seq profiles of oRGs^16^ (Fig. 1i). Using oRG-specific WTA via image segmentation, as introduced in the last results section, we detected more than 3,000 genes when profiling 6 or more oRG cells (Fig. 1i). Alternatively, we profiled small bulk ROIs in the VZ, which largely consists of ventricular radial glia (vRGs) and developing interneurons. We detected the expression of ∼4,000 genes, including many vRG and interneuron markers, from as few as 10 cells (Supp. Fig. 2c). These data demonstrate that WTA can accurately profile transcriptomes with cellular resolution, yet it provides higher sensitivity of detection when several cells are used as input material.

In the WTA assay, non-specific probe binding and incomplete transcript capture on tissue samples can lead to technical noise and obstruct accurate analysis. To estimate and reduce technical noise in WTA data, we developed the CountCorrect model (Fig. 1j) (Supp. Methods). CountCorrect estimates non-specific probe binding for each gene using a probabilistic generative model based on the negative control probes, analysis of which reveals that probe counts: (1) strongly correlate with total numbers of transcripts in a given ROI (*ρ* = 0.91, s.d. = 0.046); and (2) vary per probe (Supp. Methods). CountCorrect estimates incomplete transcript capture (sampling noise) using a Poisson model and it infers specific probe binding, which is assumed to represent true gene expression (biological signal), using non-negative matrix factorisation (Supp. Methods).

Applied to our developing brain WTA data, CountCorrect noise estimates varied per gene and distributed slightly above the mean count of negative probes (Fig. 1j). CountCorrect removed 57% of WTA probe counts on average across all genes, compared to 42% of counts that would be removed by the naive strategy of subtracting the average negative probe counts from each gene. To validate these results, we compared corrected WTA gene expression patterns to scRNA-seq measurements^16^. Reassuringly, 180 of the 200 most corrected genes (i.e. largest percentage of counts removed) in WTA data had low or no expression detected by scRNA-seq, and only 20 showed considerable expression (Supp. Fig. 2d, Supp. Table 1). In contrast, use of the naive correction strategy resulted in 55 genes with considerable expression in scRNA-seq data (Supp. Fig. 2d). Furthermore, we considered known cell type markers with moderate expression levels such as *TBR1, CX3CR1* and *CLDN5* (Supp. Fig. 2e) and found that their estimated background was lower than the average negative probe counts. Collectively, these observations suggest that CountCorrect estimates accurately reflect the underlying gene expression signal and provide a more robust approach to remove noise from WTA than naive strategies. Thus, for the remainder of this manuscript corrected counts are used, unless otherwise stated.

### Spatial mapping of cellular compartments and autism genes

Next, we sought to validate whether WTA accurately maps spatial gene expression patterns in the fetal neocortex. Initially, we examined canonical marker genes for cell types known to be localised to different cellular compartments^16,17^. WTA recapitulated the known spatial expression patterns of cell type markers across bulk ROIs along the cortical depth (Fig. 2a, Supp. Fig. 3). In the germinal zones, the markers of vRG cells (*HES1, CRYAB*) were highly enriched in the VZ, while oRG markers (*HOPX, PTN*) were enriched in the OSVZ. Intermediate progenitors (IP) are present throughout the VZ-OSVZ and their markers were elevated in these compartments relative to the cortical plate. Markers of developing interneurons (*DLX2*) that are born in the ventral telencephalon and migrate into the neocortex through the deep germinal zones were observed in the VZ-ISVZ, while markers of mature interneurons (*GABRB3*) were enriched in the cortical plate. Maturing excitatory neuron markers (*SATB2, SYT4*) were detected along their migration path from the germinal zones into the cortical plate. In the superficial cortex, markers of subplate neurons (*CRYM, TBR1*) and deep excitatory layer neurons (*SOX5, BCL11B*) were localised to their respective compartments. Finally, we observed the major classes of glial cells previously reported at this stage, oligodendrocyte precursors (*OLIG2*)^18^ and microglia (*AIF1*)^19^, as well as endothelial cells (*CLDN5*) and pericytes (*RGS5*). Collectively, these observations show that WTA accurately resolves spatial gene expression *in situ*.

**Figure 3:**
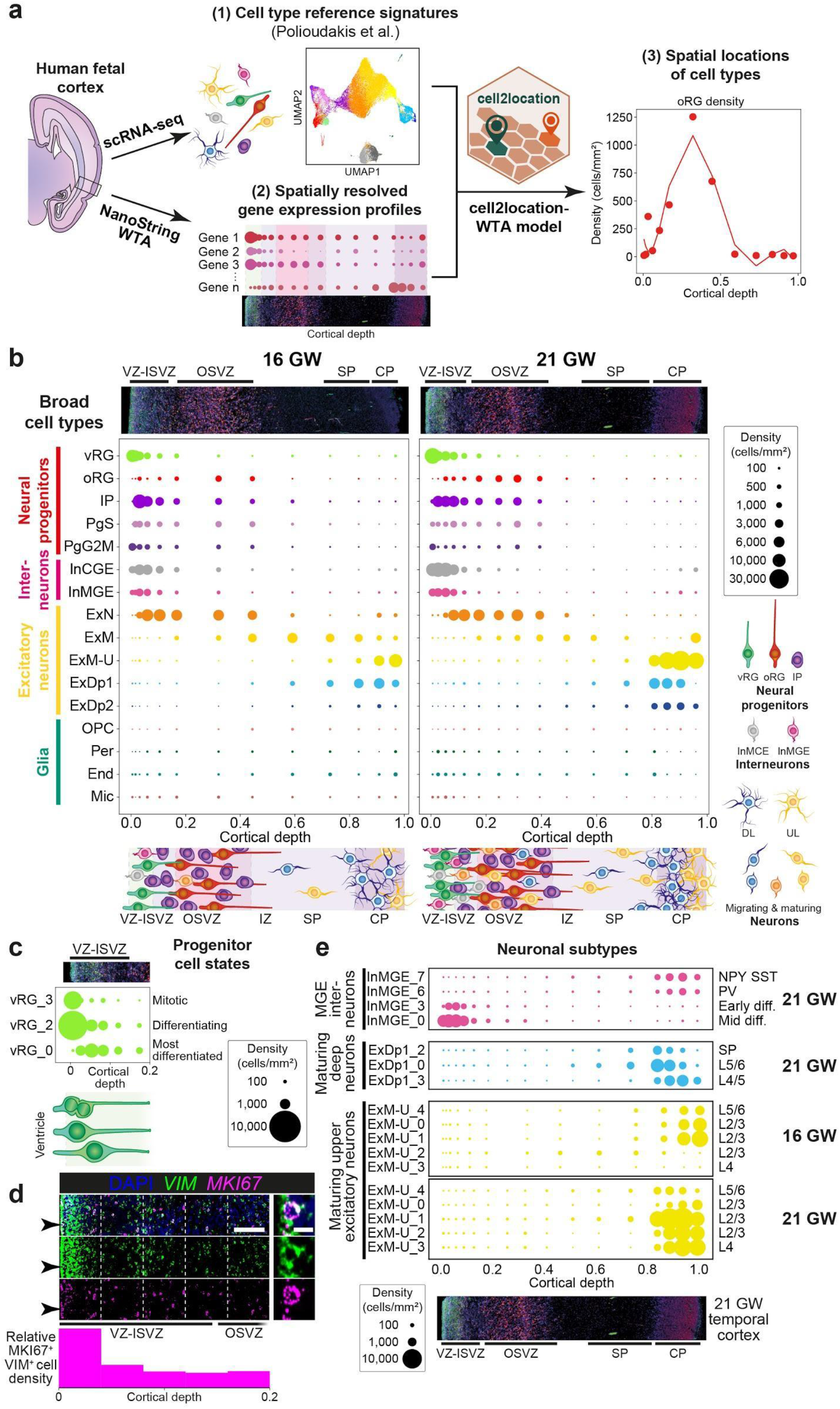
Spatial mapping of cortical cell types using integration of WTA and scRNA-seq data. **a**, cell2location-WTA workflow for spatial mapping of cell types illustrating the integration of WTA with a scRNA-seq reference dataset of the developing cortex. **b**, Spatial maps of broad cell types across the depth of the temporal cortex at 16 and 21 GW. 15,136 genes shared between the WTA and the scRNA-seq reference datasets were used for mapping, without prior marker selection. cell2location-WTA was used to estimate the absolute numbers of each cell type within each ROI, which were used to plot cell density per mm^2^ of tissue. **c**, Spatial mapping of mitotic and differentiating vRG cell states within the VZ-ISVZ. **d**, Validation of the ventricular localisation of mitotic vRGs by RNAscope smFISH. Histogram shows the quantification of MKI67+ VIM+ cells across indicated depth bins (white dashed lines) throughout the germinal zones. Scale bars, left 100 μm, right 10 μm. **e**, Mapping of neuronal subtypes reveals spatial localisation differences that reflect developing organisation of cortical layers and differentiation timing.

We then examined whether WTA can discover spatial gene expression patterns and cellular compartments in an unbiased manner. We used SpatialDE^20^ to identify and cluster spatially varying genes into patterns across cortical bulk ROIs. SpatialDE analysis successfully resolved spatial gene expression clusters from WTA data corresponding to cellular compartments and their resident cell types in the neocortex (Fig. 2b, Supp. Fig. 4). A cluster spatially enriched in the VZ (module 6) expressed marker genes for vRGs and interneurons, while OSVZ clusters (modules 1 and 3) contained markers of oRGs and IPs. Clusters mapping to the subplate and deep cortical layers (modules 5 and 7) expressed matching neuronal subtype markers (Fig. 2b). These results show WTA data enables unbiased discovery of spatially variable genes and automated mapping of tissue regions.

**Figure 4:**
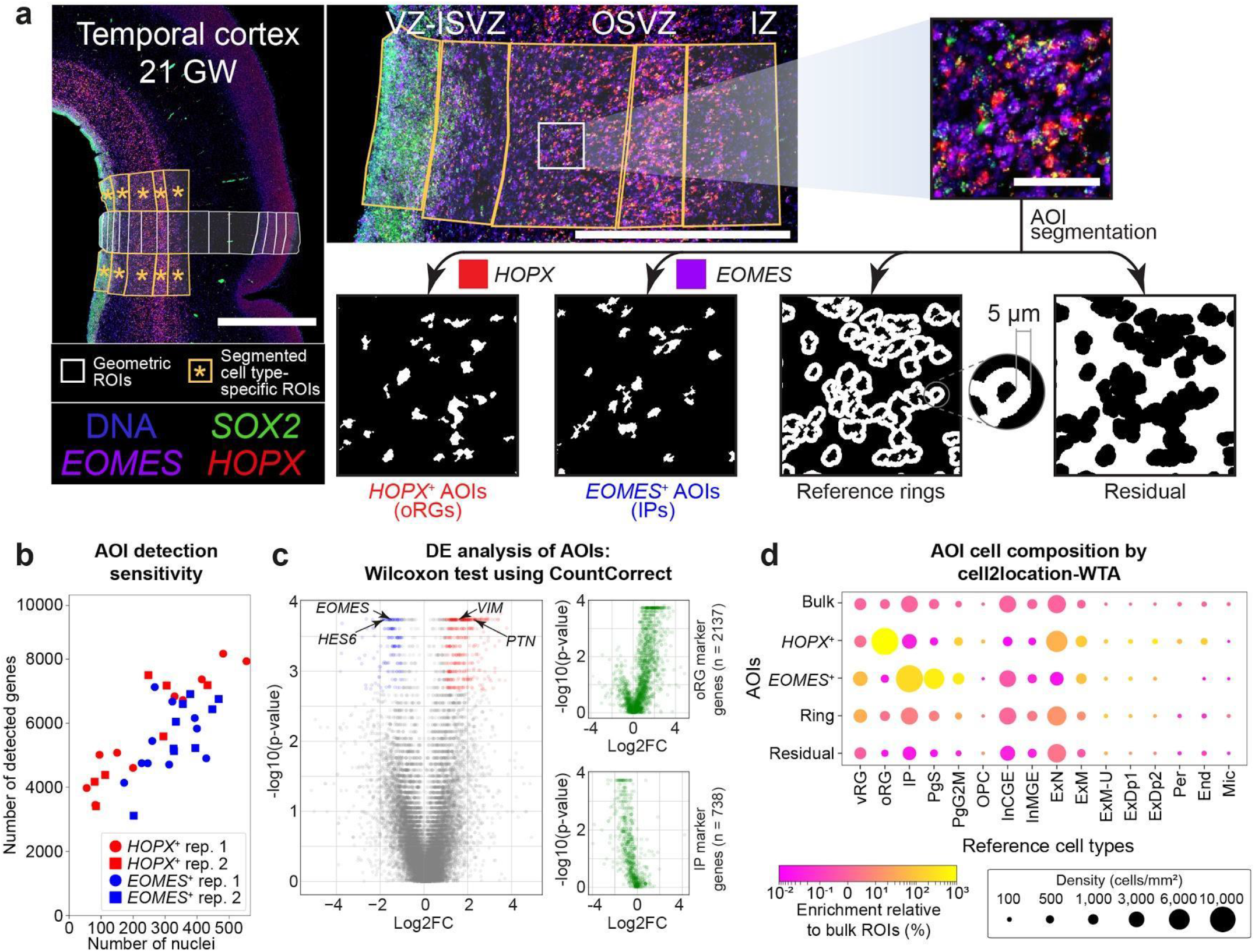
Cell type-specific WTA profiling. **a**, GeoMX custom image segmentation strategy. Bulk ROIs were segmented into cell type-specific masks on the basis of intensity thresholding of RNAscope smFISH staining. Rings of 5 μm radius surrounding masked cells provide a reference to assess efficiency and specificity of mask-oriented photocleavage. Scale bars, left 2 mm, middle 1 mm, right 100 μm. **b**, Gene expression detection sensitivity in segmented AOIs for both technical replicates of 21 GW case 1. LoD was defined as the mean plus 2 standard deviations of the negative probe counts in an AOI, calculated using log2 counts. **c**, (Left) differential expression analysis between HOPX^+^ and EOMES^+^ AOIs using the nonparametric Wilcoxon Rank Sum test on corrected counts returns 542 genes (FDR < 0.05). (Right) position of oRG and IP markers from scRNA-seq^16^ showing their enrichment in HOPX^+^ and EOMES^+^ AOIs, respectively. **d**, Analysis of AOI cell composition by cell2location-WTA indicates strong enrichment of targeted cell populations: oRGs in HOPX^+^ AOIs; and IPs, PgS and PgG2M in EOMES^+^ AOIs. Shown is the mean decomposition of ten segmented ROIs for one technical replicate of 21 GW case 1, relative to neighbouring bulk ROI data. The two most superficial HOPX+ AOIs were excluded due to the lack of HOPX+ oRGs in the VZ.

Finally, we applied WTA to spatially localise disease-related gene expression in the developing cortex. While previous studies have utilised bulk RNA-sequencing to identify the convergent expression of genes linked to autism spectrum disorder (ASD) in deep layer neurons^21^, the spatial expression pattern of most ASD genes has not been validated *in situ*. Here, we examined our WTA data to map the spatial expression of 102 ASD genes derived from a recent exome sequencing study^22^ in bulk ROIs across cortical depth. To provide ASD-specific context, we also examined the spatial expression of genes linked to brain size^23^ and IQ^24^, and a comprehensive set of neurodevelopmental disorder (NDD) genes^25^. The relative expression of each gene set (i.e. the expression mean of the gene set divided by the mean of all detected genes) showed that ASD gene expression was highly enriched in the sub-and cortical plate (Fig. 2c). While brain size and NDD genes were modestly enriched in deep and upper cortical plate, respectively, ASD genes significantly peaked in deep cortical layers with 50% higher expression than expected for a randomly chosen detected gene at both 16 and 21 GW (Fig. 2c, Supp. Fig. 5). These results provide the first high-throughput spatial validation of ASD gene convergence in deep cortical layers and demonstrate the potential of WTA for mapping disease gene expression *in situ*.

### WTA and scRNA-seq integration maps cell types at scale

Spatial mapping of resident cell types across complex tissues can offer new perspectives into cellular signalling, tissue development and function. The integration of single-cell and spatial transcriptomic data, where cell types are defined using scRNA-seq and then resolved across spatially assayed tissue locations, offers a scalable workflow to map cell types *in situ*^*26–28*^.

Here, we sought to resolve cell types in WTA data and map their spatial distribution in the developing human neocortex. We computationally integrated our WTA dataset with a scRNA-seq reference of the mid-gestation human neocortex^16^ using the cell2location model^27^. (Fig. 3a, Supp. Fig. 6a). We derived reference cell type signatures from the scRNA-seq study, using the average gene expression profiles of annotated cell type clusters^16^. We then utilised cell2location to decompose the WTA transcript counts into these reference signatures, thereby estimating the abundance of each cell type in each ROI. To account for the noise in WTA data, we incorporated CountCorrect-based background correction into the cell2location model (Supp. Methods). The extended cell2location-WTA model spatially mapped cell types with higher accuracy than the original model, providing improved enrichment of cell types to their known cellular compartments (Supp. Fig. 6b).

Initially, we examined the spatial mapping of the 16 broad cell types annotated in the scRNA-seq reference^16^. cell2location-WTA accurately mapped these broad cell types in the temporal neocortex at both 16 and 21 GW (Fig. 3b). vRG and oRG cells were mapped to the VZ and OSVZ respectively, while IPs and cycling progenitors (PgS, Pg2M) were located across the germinal zones. Interneurons (InCGE, InMGE) were enriched in the VZ-ISVZ. Excitatory neurons were also mapped accurately: newly born (ExN), maturing (ExM), maturing deep (ExDp1,2) and upper (ExM-U) layer excitatory neurons were positioned along the cortical depth according to their differentiation stage and laminar destinations. Moreover, we could detect changes in cell abundance across age (Fig. 3b), as there were higher numbers of interneurons and differentiated excitatory neurons in the cortical plate at 21 GW, consistent with increased neuronal accumulation across fetal development^12^.

Next, we used cell2location-WTA to map fine-grained cell subtypes in WTA data. This mapping resolved the spatial positions of 78 progenitor and neuronal subtypes annotated in the scRNA-seq reference^16^, assigning them to expected cellular compartments (Supp. Fig. 7). Amongst progenitors, mitotic and early differentiated vRG subtypes were mapped to the ventricular surface and adjacent areas in VZ, respectively (Fig. 3c), consistent with previous observations^29^ and smFISH validation (Fig. 3d). In contrast, more differentiated vRGs expressing neuronal markers^16^ were positioned away from the ventricle (Fig. 3c). Amongst neuronal subtypes, differentiating interneurons (InMGE_0,3) were located in the VZ-ISVZ while more mature PV and SST interneurons (InMGE_6,7) were mapped to the cortical plate (Fig. 3e). Deep layer excitatory neuronal subtypes annotated as subplate, layer 5/6 and layer 4/5 populations were also accurately mapped across cortical depth (Fig. 3e).

Finally, we could also resolve the migration order of fine neuronal subtypes into the cortical plate. While it is well known that deep and upper layer excitatory neurons are sequentially generated over development and that they subsequently migrate into the cortical plate^30^, the precise migration order of fine subtypes within each lamina is less well characterised. We found that amongst maturing excitatory (ExM-U) neurons, layer 5/6 (ExM-U_4) and 2/3 (ExM-U_0,1) subtypes were observed at the cortical plate at 16 GW, while two distinct layer 2-4 subtypes (ExM-U_2,3) were observed only later at 21 GW (Fig. 3e), indicating their sequential migration order. Taken together, these results demonstrate that the integration of WTA and scRNA-seq can spatially map cell types at scale and resolve the locations of closely related subtypes *in situ*.

### Cell type-specific WTA profiling

The DSP technology can target specific cell types via image segmentation of fluorescent visualisation markers and spatially precise photocleavage of probe barcodes^8^. Here, we tested whether WTA can accurately capture cell type-specific transcriptomes *in situ* by profiling two different fetal neocortical progenitor populations, outer radial glial stem cells (oRGs) and intermediate progenitors (IPs). After WTA probe hybridisation and visualisation marker imaging, we used image segmentation to identify *HOPX*^+^ and *EOMES*^+^ areas of interest (AOIs) corresponding to the cell bodies of oRGs and IPs, respectively, in segmental ROIs across the 16 and 21 GW germinal zones (Fig. 4, Supp. Fig. 8). We sequentially collected WTA probe tags from the *HOPX*^+^ and *EOMES*^+^ AOIs within each ROI, as well as “reference rings’’ surrounding the segmented cells and the residual area to assess the accuracy of cell type profiling (Fig. 4a). To facilitate comparison of cell type-specific and bulk WTA profiling, we placed segmented ROIs adjacent to those used for bulk profiling earlier in our study (Fig. 4a, Supp. Fig. 8a-b).

The cell type-specific AOIs contained an estimated 100 to 400 nuclei each, where we detected the expression of 4,000-8,000 genes per AOI using a negative probe count based threshold (Fig. 4b). We employed the non-parametric Wilcoxon test to identify 542 differentially expressed genes between *HOPX*^*+*^ and *EOMES*^*+*^ AOIs processed by CountCorrect (Fig. 4c). In contrast, the same test applied to *HOPX*^*+*^ and *EOMES*^*+*^ clusters from the scRNA-seq data^16^ revealed 4,155 differentially expressed genes. Comparison of the two sets revealed 343 genes in common, including many known cell type markers (Fig. 4c). Closer inspection of the genes that were identified as differentially expressed in scRNA-seq but not WTA revealed their enrichment in the relevant cell type specific AOI in WTA data (Fig. 4c, small panels). Finally, we identified fewer genes when we applied the same differential expression test to uncorrected WTA data (175 genes) and data corrected by subtracting the mean expression of negative probes (414 genes) (Supp. Fig. 9). Taken together, these results suggest that cell type profiling by WTA is consistent with scRNA-seq and is improved by CountCorrect.

To further assess the cell-type specificity of WTA profiling, we quantified the cell type composition of our AOIs (*HOPX*^+^, *EOMES*^+^, reference rings, residual) and bulk ROIs using cell2location-WTA (Fig. 4d, Supp. Fig. 8c). Relative to bulk ROIs, *HOPX*^+^ AOIs were highly enriched for oRGs (11.6-fold), while *EOMES*^+^ AOIs contained elevated numbers of IPs (3.3-fold enrichment) and the related cycling progenitor subtypes PgS and PgG2M (6.1-fold, 5.4-fold). Consistently, the targeted cell types were depleted from the corresponding opposite segmental AOIs as well as the reference rings and the residual AOI space. While *HOPX*^+^ and *EOMES*^+^ AOIs were enriched to a lesser degree for maturing excitatory neurons and vRGs respectively (Fig. 4d), likely due to the challenging nature of segmentation in the densely packed germinal zones, the high level of enrichment for targeted cell types demonstrates that WTA can be used to capture transcriptomes from specific cell populations *in situ*.

Finally, to assess whether WTA can be used to explore cell-cell communication, we examined whether it can validate the extracellular receptor-ligand interactions involving oRGs and IPs extracted from scRNA-seq data by the CellPhoneDB algorithm^31^. The majority of receptors and ligands expressed in oRG and IP clusters in scRNA-seq data^16^ were also detected in their respective WTA AOIs (Supp. Fig. 10a, Supp. Table 3). Focusing on oRGs, we found that the majority (113/124) of oRG-specific interactions identified in scRNA-seq data were also detected in WTA data (Supp. Fig. 10b), where the oRG receptors/ligands were expressed in *HOPX*^+^ AOIs and their interaction partners were expressed in *HOPX*^-^ “background” AOIs in a complementary manner. These included previously documented interactions such as LIF-LIFR, PTN-PTPRZ1 and FGF2-FGFR2^17^. Finally, we were able to validate interactions such as that between oRG-expressed HBEGF and interneuron-expressed ERBB4 with RNAscope (Supp. Fig. 11a-b), which had not been mapped *in situ* previously. Overall, this shows that WTA can be used to comprehensively screen cell-cell interactions.

## Discussion

Transcriptome-wide spatial profiling promises valuable insights into tissue architecture in health and disease. Here, we presented the NanoString WTA technology that can spatially resolve the expression of over 18,000 genes on fixed tissue samples. We demonstrated that WTA enables unbiased *in situ* identification of spatial gene expression patterns, including that of disease-related genes and cell-cell signalling molecules. Furthermore, WTA can be integrated with single-cell transcriptomics to comprehensively map fine cell types across complex tissues. Hence, the assay is not limited to the spatial validation of predefined cell type or disease biomarker panels and WTA can be employed as a discovery tool for cell atlasing and pathology.

We demonstrated diverse spatial transcriptomic applications of WTA, ranging from profiling large tissue compartments to single cells. While most spatial RNA-seq methods require profiling whole tissue sections in a gridded manner, the DSP methodology allows WTA to target tissue areas of diverse shapes or sizes or specific groups of cells. Here, we illustrated WTA profiling of segmental tissue regions to map tissue atlases at a coarse spatial resolution. We also demonstrated that WTA can capture cell type specific transcriptomes. In particular, cell type specific WTA can be readily applied as a discovery tool for pathology, where many cellular targets such as immune or tumor subtypes can be reliably identified by few biomarkers. This approach can also selectively profile the full transcriptomes of disease-relevant cell types at a lower cost than grid-sequencing entire tissue sections.

We validated WTA in the developing human brain, creating the first spatial transcriptomic map of the fetal temporal cortex. There are several promising future directions for WTA profiling in developmental neuroscience. WTA can be scaled to profile multiple brain regions throughout embryonic and postnatal life, creating a spatial atlas of human brain development that could be readily integrated with scRNA-seq datasets^15,32^. Our spatial risk gene enrichment approach can also be extended accordingly to identify regional neural circuits or developmental stages relevant to the aetiology of neurodevelopmental disorders such as autism. Furthermore, WTA can be applied to archival tissue samples to identify cell type-specific aberrations in neurodevelopmental disorders.

Similar to other genomic technologies, WTA experiments contain measurement noise. To de-noise WTA data, we formalised the generative process of WTA experiments in a statistical model called CountCorrect. This approach enabled us to control for these noise sources in downstream analyses and improve the identification of differential gene expressed genes and cell type decomposition. We envision that the statistical approach in CountCorrect could also be used in other high-throughput probe based technologies, such as in situ sequencing^3^.

To accurately map cell types in WTA data, we incorporated CountCorrect into the cell2location spatial mapping algorithm^27^. The resulting cell2location-WTA model provides a scalable workflow to integrate scRNA-seq and WTA data and estimate the abundance of individual cell types in bulk or segmented WTA datasets. Hence, our approach can be applied to coarse spatial resolution WTA data, such as that from large tissue compartments, to map tissue atlases with higher cell type resolution.

Our cell2location-WTA workflow offers several benefits as demonstrated in our human brain data. First, it provides a Bayesian framework to accurately map complex tissues with dozens of cell types and allows to distinguish fine-grained subtypes with subtle transcriptional differences. Second, it incorporates CountCorrect to accurately model the measurement sensitivity of WTA experiments and improve cell type mapping. Third, in addition to estimating cell proportions, it enables us to estimate absolute cell abundance. Finally, cell2location-WTA is computationally efficient and allows use of the full transcriptome WTA data without prior gene selection for spatial mapping. Hence, we expect cell2location-WTA will be applied to comprehensive tissue atlasing as well as clinical research where it can spatially resolve cellular changes in pathological conditions.

We envision future developments of CountCorrect to further improve noise removal and cell type mapping in WTA data. We also expect more robust models for analysing cell type specific WTA data that can yield more differentially expressed genes akin to scRNA-seq observations. In addition, as more WTA datasets become available, batch correction approaches will be required to combine data for unified analysis.

Importantly, future work can integrate the image and transcriptome data from WTA measurements to correlate tissue and cell morphology with gene expression profiles. This can provide valuable insights into tissue development and function, and help identify molecular processes that organise tissue architecture. Furthermore, it could identify morphological correlates of molecular disease signatures and thus generate imaging biomarkers to inform about a patient’s disease state from imaging data alone.

In summary, we presented WTA as a versatile transcriptome-wide discovery tool for cell atlasing and fixed tissue spatial transcriptomics. We expect WTA to be widely applicable to tissue mapping and analysis of clinical tissue samples.

## Methods

### Human tissue

Formalin-fixed paraffin-embedded (FFPE) blocks of second trimester human fetal brain were obtained from the Human Developmental Biology Resource, Newcastle, UK (REC 18/NE/0290).

### RNAscope *in situ* hybridisation

Sections were cut from FFPE blocks at a thickness of 5 μm using a Leica microtome, placed onto SuperFrost Plus slides (VWR), and baked overnight at 55°C to dry and ensure adhesion. Tissue sections were then processed using a Leica BOND RX to automate staining with the RNAscope Multiplex Fluorescent Reagent Kit v2 Assay (Advanced Cell Diagnostics, Bio-Techne), according to the manufacturers’ instructions. Automated processing included baking at 60°C for 30 minutes and dewaxing, as well as heat-induced epitope retrieval at 95°C for 15 minutes in buffer ER2 and digestion with Protease III for 15 minutes.

For visualising markers prior to NanoString GeoMX profiling, 3-plex RNAscope was developed using tyramide signal amplification with Opal 570, Opal 620, and Opal 690 dyes (Akoya Biosciences). No nuclear stain was applied at this stage.

For validation staining, 3-plex or 4-plex RNAscope stains were developed using Opal 520, Opal 570, and Opal 650 dyes (Akoya Biosciences), as well as TSA-biotin and streptavidin-conjugated Atto 425 (Sigma). Nuclei were counterstained with DAPI at 167 ng/ml.

### Confocal imaging

Imaging of validation RNAscope-stained slides was performed using a Perkin Elmer Opera Phenix High-Content Screening System, in confocal mode with 1 μm z-step size, using 20× (NA 0.16, 0.299 μm/pixel) or 40× (NA 1.1, 0.149 μm/pixel) water-immersion objectives. Channels: DAPI (excitation 375 nm, emission 435-480 nm), Atto 425 (ex. 425 nm, em. 463-501 nm), Opal 520 (ex. 488 nm, em. 500-550 nm), Opal 570 (ex. 561 nm, em. 70-630 nm), Opal 650 (ex. 640 nm, em. 650-760 nm).

### GeoMX Whole Transcriptome Atlas slide preparation

In addition to the use of RNase-free reagents, surfaces, equipment, and staining containers were cleaned using RNase AWAY Surface Decontaminant (Thermo Scientific) throughout slide processing.

Following RNAscope staining, slides were processed according to the NanoString GeoMX RNA assay protocol (MAN-10087-02). Sections were briefly rinsed in nuclease-free water, and then post-fixed in 10% neutral-buffered formalin for 5 minutes. Fixation was quenched by incubation in 0.1 M glycine, 0.1 M Tris, twice for 5 minutes each, followed by 5 minutes washing in PBS, whereafter probes were applied immediately, again according to the assay protocol.

The Whole Transcriptome Atlas (WTA) probe reagent (For Research Use Only) was diluted in pre-equilibrated buffer R to a final probe concentration of 4 nM and added to each slide, which was covered with a Hybrislip cover (Grace Bio-Labs) and incubated for 15 hours at 37°C in a HybEZ II System, humidified with 2× SSC (saline-sodium citrate) buffer. The following day, slides were de-coverslipped by brief rinsing in 2× SSC with 0.05% Tween-20, and then washed twice for 25 minutes each in 2× SSC, 50% formamide at 37°C, and twice for 5 minutes each in 2× SSC at room temperature.

Following washing, slides were counterstained with the DNA dye SYTO13. A SYTO13 stock (5 μM) was clarified by centrifugation at 13,000 g for 2 minutes, and then diluted to 500 nM in buffer W prior to staining in the dark for 30 minutes. Finally, slides were washed twice for 3 minutes each in 2× SSC buffer.

### GeoMX ROI collection and segmentation

Slides were covered with buffer S and loaded into the GeoMX DSP instrument. Regions of interest (ROIs) were selected using the polygon tool; a subset of annotated ROIs were then exported for cell segmentation.

ROIs were segmented into AOIs (areas of interest) based upon staining for the intermediate progenitor marker *EOMES* and the outer radial glia marker *HOPX*, using a custom Rare Cell ImageJ script applied independently to the two respective image channels. The resulting masks were combined and then used to produce the reference ring and residual masks using a custom Contour ImageJ script. At the time of experimentation, the script did not support segmentation of multiple populations. As a result, the independent population masks were combined in Adobe Photoshop using Exclude and Subtract layering to remove any minor regions of overlap, which would prevent import into the GeoMX DSP.

Following collection of sequencing tag-containing aspirates, wells were dried either at room temperature overnight or at 65°C for 45 mins in a heating block, and then re-suspended in 10 μl of nuclease-free water (Ambion), in order to minimise any differences due to ambient evaporation.

### Library preparation and sequencing

ROI-derived oligos were each uniquely dual-indexed using the i5 x i7 system (Illumina). A 4 μl aliquot of each re-suspended ROI aspirate containing the photocleaved oligos was amplified in a PCR reaction containing 1 μM i5 and i7 primers and 1× NSTG PCR Master Mix. UDG digestion was carried out at 37°C for 30 min, and then deactivated at 50°C prior to denaturation at 95°C for 3 minutes, and 18 cycles of amplification: 95°C for 15 seconds, 65°C for 1 minute, 68°C for 30 seconds. Final extension was conducted at 68°C for 5 minutes.

Prior to purification, PCR reactions were pooled into two mixtures, separating ‘large’ and ‘small’ ROIs. The large pool included geometric ROIs (excluding the sensitivity-assaying series featured in Supp. Fig. 2c) and the residual area from segmented ROIs. The small pool included all *EOMES*^+^ and *HOPX*^+^ segmented populations, associated reference rings, and the small geometric ROIs omitted from the large pool. The two pools of PCR reactions were each purified with two rounds of AMPure XP beads (Beckman Coulter) at 1.2× sample volume of beads.

Large-ROI and small-ROI libraries were quantified using an Agilent 2100 Bioanalyzer and High Sensitivity DNA Kit; integration of the 161 bp amplicon peak indicated concentrations of 13 nM and 21 nM, respectively. The libraries were pooled in a biased manner designed to target the large ROI pool with two-thirds of reads, and sequenced with 50PE reads across both lanes of an Illumina NovaSeq 6000 S2 flow cell at a concentration of 400 pM, with 5% PhiX spike-in, yielding 2.4 billion reads.

### NanoString GeoMX data processing

DSP sequencing data were processed with the GeoMx NGS Pipeline (DND). After sequencing, reads were trimmed, merged, and aligned to a list of indexing oligos to identify the source probe. The unique molecular identifier (UMI) region of each read was used to remove PCR duplicates and duplicate reads, thus converting reads into digital counts.

### Number of detected genes

The limit of detection (LoD) in an ROI was defined based on the mean and standard deviation (s.d.) of log2-normalised negative probe counts. On the log scale the calculation is: LoD = mean + (2 × s.d.). For the raw negative probe counts this corresponds to: LoD = mean × s.d.^2^. We used this LoD threshold to produce Figures 1f,i. To count the total number of detected genes in cortical tissue across geometric ROIs at 21 GW, we used all geometric ROIs from biological case 1, as well as 16 cortical ROIs from case 2. A gene was counted as detected only if it exceeded the LoD in at least 3 of those ROIs. For the interaction analysis in AOIs and Supp. Fig. 10a, we similarly counted a gene as detected only if it exceeded the LoD threshold in at least 3 AOIs out of all *HOPX*^+^ and *EOMES*^+^ AOIs.

A gene was counted as detected in the reference scRNA-seq dataset, if its average UMI counts exceeded 0.02 in a cell type cluster. We used this threshold to count the total number of detected genes in the dataset, as well as the receptor/ligand genes detected for Supp. Fig. 10a.

### Reproducibility

To calculate technical and biological reproducibility in ROIs for Fig. 1g-h, we chose the most superficial geometric ROI in the cortical plate on all slides. The Pearson correlation values were calculated on log2 counts-per-million (CPM). To calculate the reproducibility of AOIs in Supp. Fig. 8d, we chose the second to outermost AOI on the chosen trajectory, at approximately cortical depth 0.3. Again, Pearson correlation values were calculated on log2 CPM.

### Corrected counts, Standardised counts (z-score), relative expression

Excluding the results in Fig. 1 and Supp. Fig. 8d, all analysis was carried out using the corrected counts from CountCorrect as a starting point (see Supp. Methods).

We calculated standardised counts/z-scores for Fig. 2a-b using the z-score function in the scipy.stats package. For each calculation we included samples from the same ventricular zone to cortical plate trajectory. The basis for the calculation was log2 counts-per-million. We took the mean of the standardised counts for Supp. Fig. 5a.

Relative expression for Fig. 2c and Supp. Fig. 5a was calculated as the ratio between mean CPM of genes in the relevant gene set and mean CPM of all detected genes on the chosen strip of 21 GW AOIs on both technical replicates.

### Spatially varying genes and gene clusters

We used the NaiveDE and SpatialDE python package to obtain significantly varying genes and cluster them into patterns. Following the spatialDE workflow, ROI counts were first stabilised using the NaiveDE “stabilize” function, followed by normalisation for the total number of counts in an ROI using the NaiveDE “regress_out” function. Finally, spatially varying genes were obtained using the SpatialDE “run” function. This identified 4,303 spatially varying genes with a q-value below 0.05. We chose to cluster all genes with at least moderate evidence of spatial variation (16,161 genes with SpatialDE qval < 0.5) into patterns using SpatialDE “spatial_patterns” function, with the length scale parameter “l” set to 0.9. This cutoff of qval < 0.5 for clustering, was taken from the main SpatialDE tutorial on github. We set the “expected number of patterns parameter” to 3, 5, 7, 10, 12 and 15 patterns in exploratory analysis. After inspecting the results, 12 patterns were chosen as the most detailed and representative of tissue biology based on our prior knowledge of cell type numbers and positioning in the developing cortex. Two of the 12 patterns were removed from further analysis. The first one contained 0 genes, the second one only contained technical outlier gene probes with very low counts in all but one ROI. Fig. 2b shows the 70 genes with highest SpatialDE membership score for each pattern in light grey, as well as common *a priori* chosen cell type marker genes.

### Cell type mapping

We used our adapted version of the cell2location-WTA method (details in Supp. Methods and the cell2location manuscript^27^) to obtain cell number estimates for each ROI/AOI in Figures 2 and 4d, as well as Supp. Figures 6, 7 and 8c. We constructed cell type reference expression profiles by averaging the counts in each cell type cluster in the reference study^16,27^, resulting in 16 profiles for the main types and 78 profiles for the subtypes. We supplied raw (i.e. unnormalised, uncorrected) Nanostring WTA counts to the algorithm. Both the reference expression profiles and the WTA data were subset to 15,136 common genes. Further parameter settings for the cell2location algorithm were as follows:

– Training iterations: 50,000
– Learning rate: 0.001
– Prior on cells per location: Mean for each ROI was specified as the nuclei counts estimated by the Nanostring software for each ROI, based on DAPI stains on the image. Standard deviation was set to 10% of the mean (CV, representing prior strength, of 0.1).
– Prior on cell types per location: Mean of 6. Default CV of 1.
– Cell type combinations per location: Mean of 5. Default CV of 1.
– Prior on difference between technologies: Mean of 0.5. SD of 0.125. CV of 0.25 for both.

### Differential expression analysis between *HOPX*^+^ and *EOMES*^+^ AOIs

A reference list of differentially expressed genes was compiled by performing a Wilcoxon rank sum test between relevant cell clusters in the single-cell RNAseq reference dataset^16^ and selecting genes with an FDR < 0.05 and log2 fold-change greater than 1. Cell clusters corresponding to the *HOPX*^+^ AOIs were oRGs, as well as two *HOPX*-expressing progenitor subclusters, denoted as ‘PgG2M_0’ and ‘PgS_4’ in the original study. Cell clusters corresponding to *EOMES*^+^ AOIs were IPs, as well as all remaining PgG2M and PgS subclusters, since they express *EOMES* in the scRNA-seq data.

### Cell-cell interaction analysis

CellphoneDB^31^ was used as a database for ligand-receptor interactions. Ligands and receptors were classified as detected/undetected as explained in the previous section on “Number of detected genes”. Importantly, for multi-subunit ligands or receptors we required all corresponding transcripts to be detected to count the ligand or receptor as overall detected. We ran the CellphoneDB algorithm on the reference single-cell RNAseq dataset to extract putative oRG interactions. Interactions with a *p*-value below 0.05 were counted as significant, as this is suggested in the CellPhoneDB algorithm by default. An interaction was counted as detected in WTA data if both the oRG ligand/receptor was detected in *HOPX*^+^ AOIs and the ligand/receptor from other cell types in the background AOIs (i.e. residual and *EOMES*^+^ AOIs).

### Image analysis

*ERBB4* smFISH signal (Supp. Fig 11b) was quantified across a strip of the ventricle-cortical depth using ilastik^33^. A pixel classifier was trained to extract the borders of the tissue section at the ventricular zone (VZ) and the cortical plate (CP); a second pixel classifier was trained to identify *ERBB4* RNAscope signal puncta. For all detected signals, distances from the signal centroid to the nearest VZ edge and nearest CP edge were measured, and the relative depth of the signal on the VZ-CP trajectory was calculated. Signals were binned according to this relative depth into 30 bins across the VZ-CP trajectory; only the first 12 representing depth 0 to 0.4 are presented.

## Data availability

WTA experiment data and sample metadata are presented as Supp. Tables 4 and 5, respectively.

## Code availability

CountCorrect python package: https://github.com/BayraktarLab/CountCorrect CountCorrect-Model in pymc3:

https://github.com/BayraktarLab/CountCorrect/tree/main/countcorrect/ProbeCounts_GeneralModel.py

Notebook for CountCorrect Model: https://github.com/BayraktarLab/CountCorrect/blob/main/BackgroundCorrection.ipynbcell2location-WTA Model:

https://github.com/BayraktarLab/cell2location/blob/master/cell2location/models/LocationModelWTA.py

Notebook for cell2location-WTA Model: https://github.com/BayraktarLab/cell2location/blob/master/docs/notebooks/cell2loation_for_NanostringWTA.ipynb

## Acknowledgements

We gratefully acknowledge: Liz Tuck for sectioning; Michael Quail and Tarryn Porter for assistance with library preparation and sequencing coordination; Mathias Holpert and Trieu My Van for training and assistance on the NanoString GeoMX DSP instrument; Arya Bahrami for the ImageJ segmentation scripts; and Jimmy Lee for help testing the CountCorrect software. Human brain samples were obtained from the MRC-Wellcome Trust Human Developmental Biology Resource.

## Author contributions

K.R., A.A. and O.A.B. conceived of the study. K.R. performed all WTA and smFISH experiments. R.F. and A.A. performed AOI image segmentation. R.F. performed library preparation and WTA decoding. A.A. performed all data analysis, developed the CountCorrect model with feedback from M.H. and the cell2location-WTA model with contributions from V.K. T.L. performed smFISH image processing. M.R. and J.M.B. participated in experimental design. K.R., A.A. and O.A.B. wrote the manuscript with feedback from all authors.

## Supplementary data

### Supplementary figures S1 to S11

***SupplementaryTable1-Top200-MostCorrected-Genes*** *shows the top 200 genes with the highest fraction of counts removed for 21GW biological case 1, technical replicate 1, according to the CountCorrect model (sheet 1) and the naive strategy (sheet 2). Included are also the total and mean WTA counts on the section for each gene*.

***SupplementaryTable2_SpatialDE*** *shows output of spatialDE spatially variable gene estimation ordered by q-value (sheet 1) for 21 GW biological case 1, technical replicate1. Sheet 2 shows classification of genes into each of the 12 modules*.

***SupplementaryTable3_Interactions*** *shows all cellPhoneDB receptors and ligands with their corresponding detection in the WTA AOIs or scRNAseq clusters (sheets 1 and 2). The oRG specific interactions detected in WTA are on sheet 3, with undetected interactions on sheet 4 and interactions with genes not included in the WTA panel in sheet 5*.

***SupplementaryTable4_WTA-Experiment_ProbeCounts*** *contains the UMI counts for each probe. It contains both gene probes and negative control probes (called NegProbe-WTX) for all samples in this study*.

***SupplementaryTable5_WTA-Experiment_Metadata*** *contains the metadata for each ROI/AOI profiled with WTA. Rows in this file are in the same order as columns in Supp. Table 4 and can also be matched by comparing the row and column identifiers respectively*.

## Supplementary Figures

**Supp. Figure 1:**
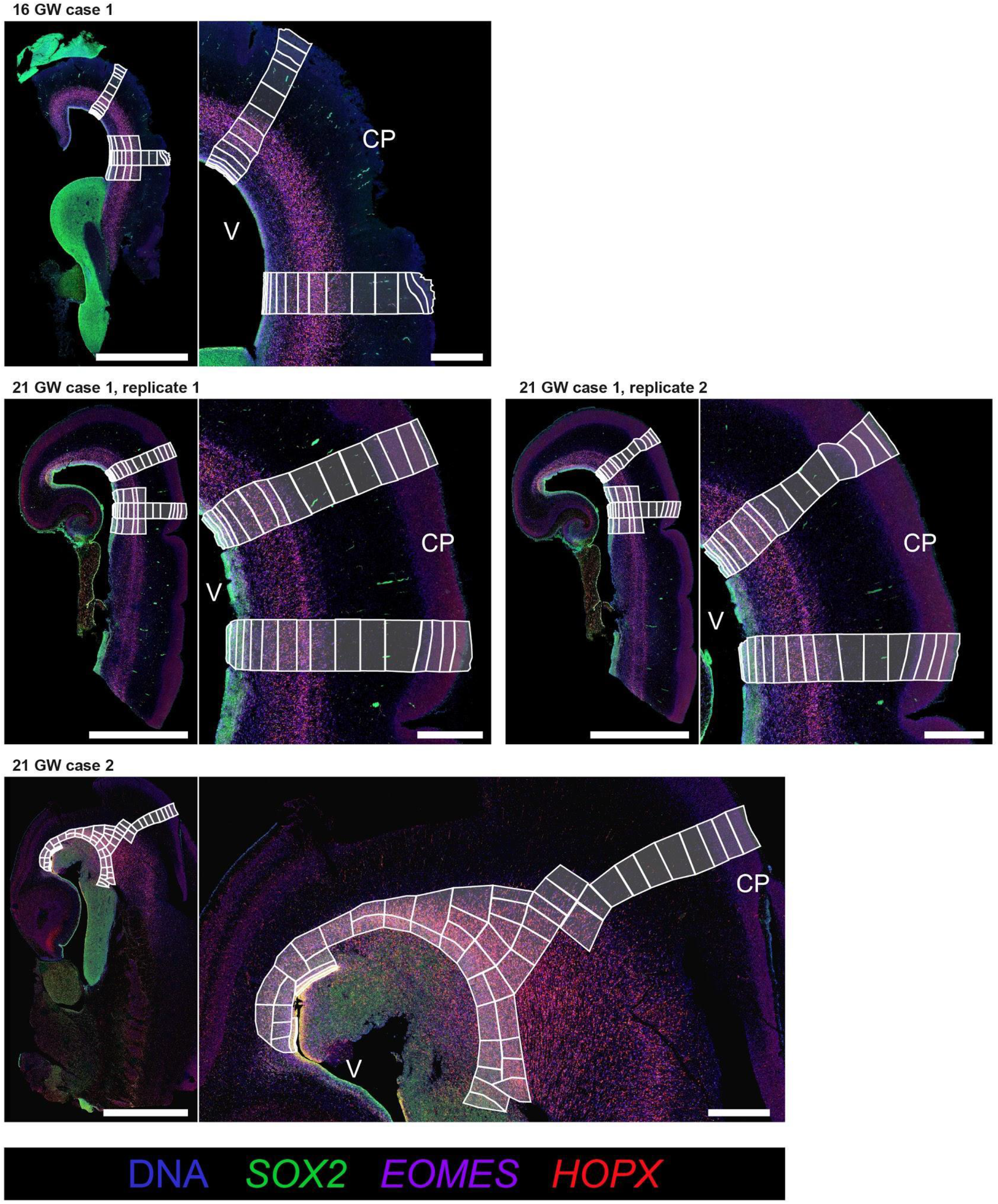
Experimental design showing all assayed fetal brain samples. Experimental design shown for all three cases, including two technical replicates (adjacent tissue sections) of 21 GW case 1. Left of each panel, fetal brain sections stained for canonical cell type markers with 3-plex RNAscope smFISH. Scale bar, 5 mm. Right, magnified view. Scale bar, 1 mm. white boxes represent individual NanoString ROIs. V, ventricle; CP, cortical plate.

**Supp. Figure 2:**
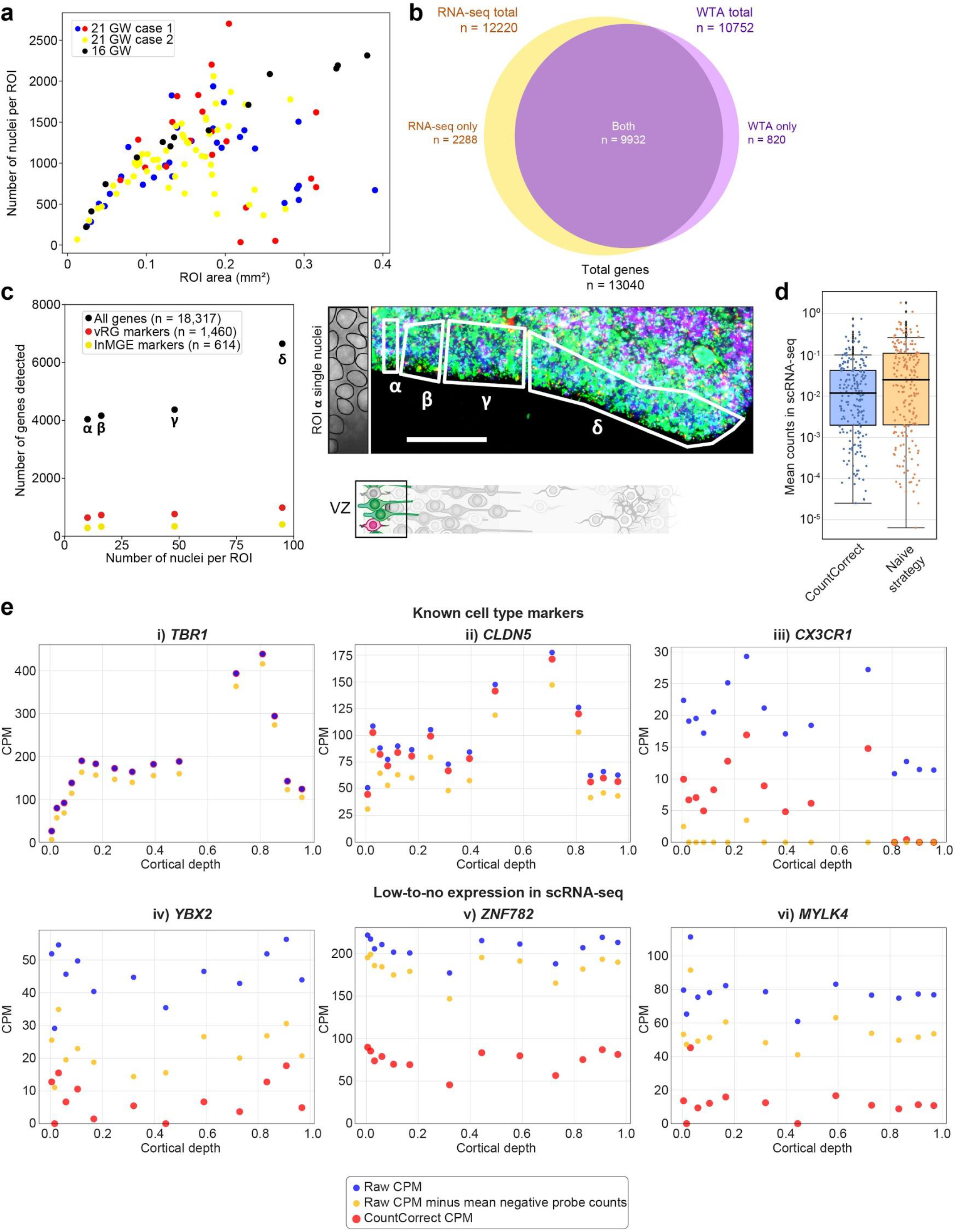
WTA detection sensitivity and de-noising by CountCorrect. **a**, Comparison of ROI size with captured nuclei number across geometric ROIs. Shown are all three cases, including two technical replicates of 21 GW case 1. **b**, Genes detected by WTA versus genes detected by scRNA-seq of the mid-gestational cortex^16^. **c**, Gene expression detection sensitivity in small geometric ROIs of varying sizes in the ventricular zone. Around 4,000 genes were detected above a negative probe count threshold from as few as 10 cells, including many markers of the dominant cell types in the region, vRGs and INs. Reference cell type markers were extracted from Polioudakis et al.^16^ using Seurat find markers. **d**, scRNA-seq average expression (across 16 main cell types) of 200 most highly corrected genes by CountCorrect and the naive strategy. The CountCorrect gene set is more lowly expressed on average and includes fewer genes with considerable expression (above 0.1 counts). **e**, Example correction patterns for (Top) known cell type markers and (Bottom) putative unexpressed genes (at most 0.01 counts in at most one cell type in scRNA-seq). CountCorrect corrections are more conservative for known cell type markers and more stringent for putative unexpressed genes. TBR1 is a subplate/deep layer neuron marker, CLDN5 marks endothelial cells, CX3CR1 corresponds to microglia.

**Supp. Figure 3:**
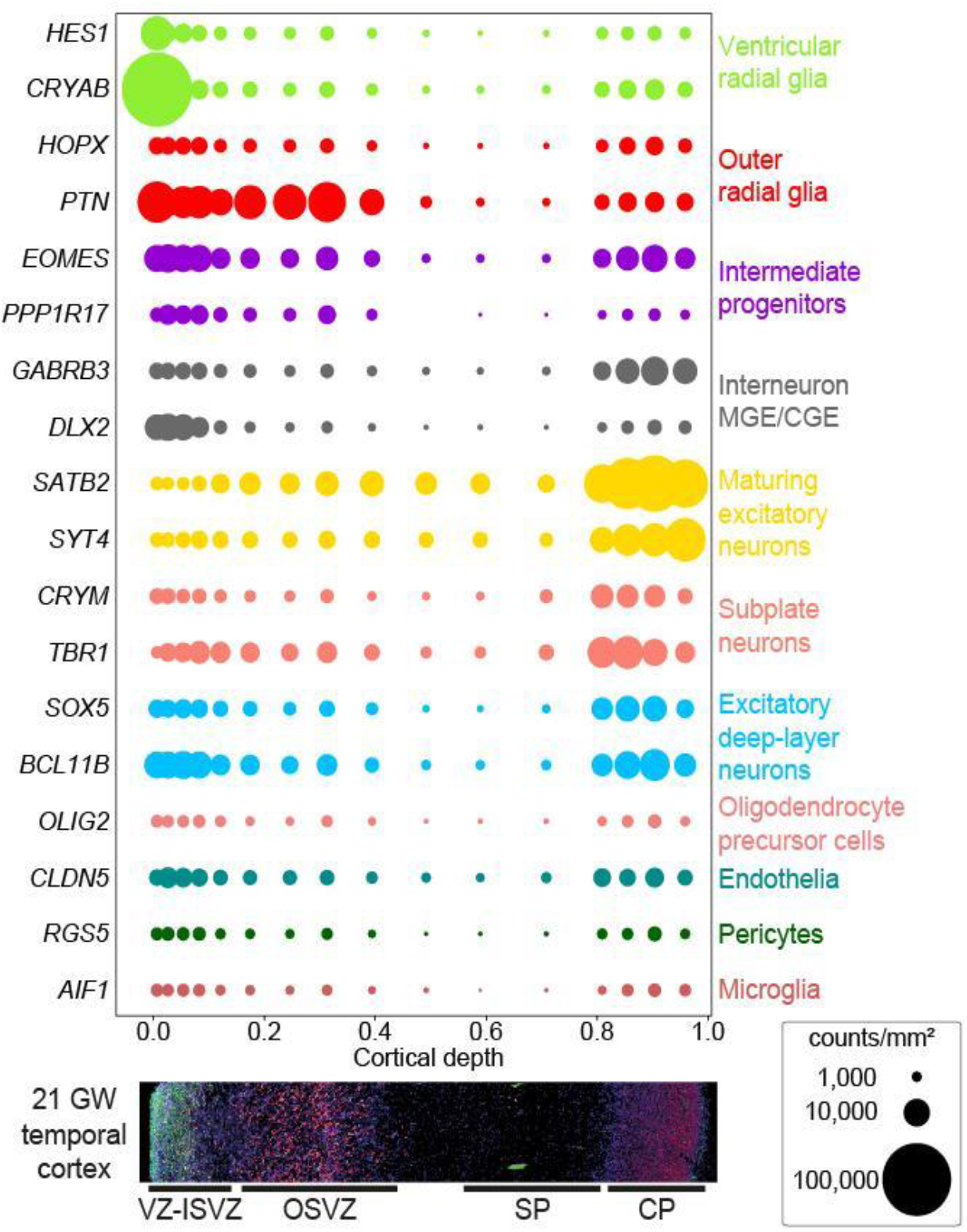
Marker gene expression across bulk WTA ROIs. Expression of canonical cell type markers across one strip of ROIs, shown as counts per mm^2^ for one technical replicate of 21 GW case 1.

**Supp. Figure 4:**
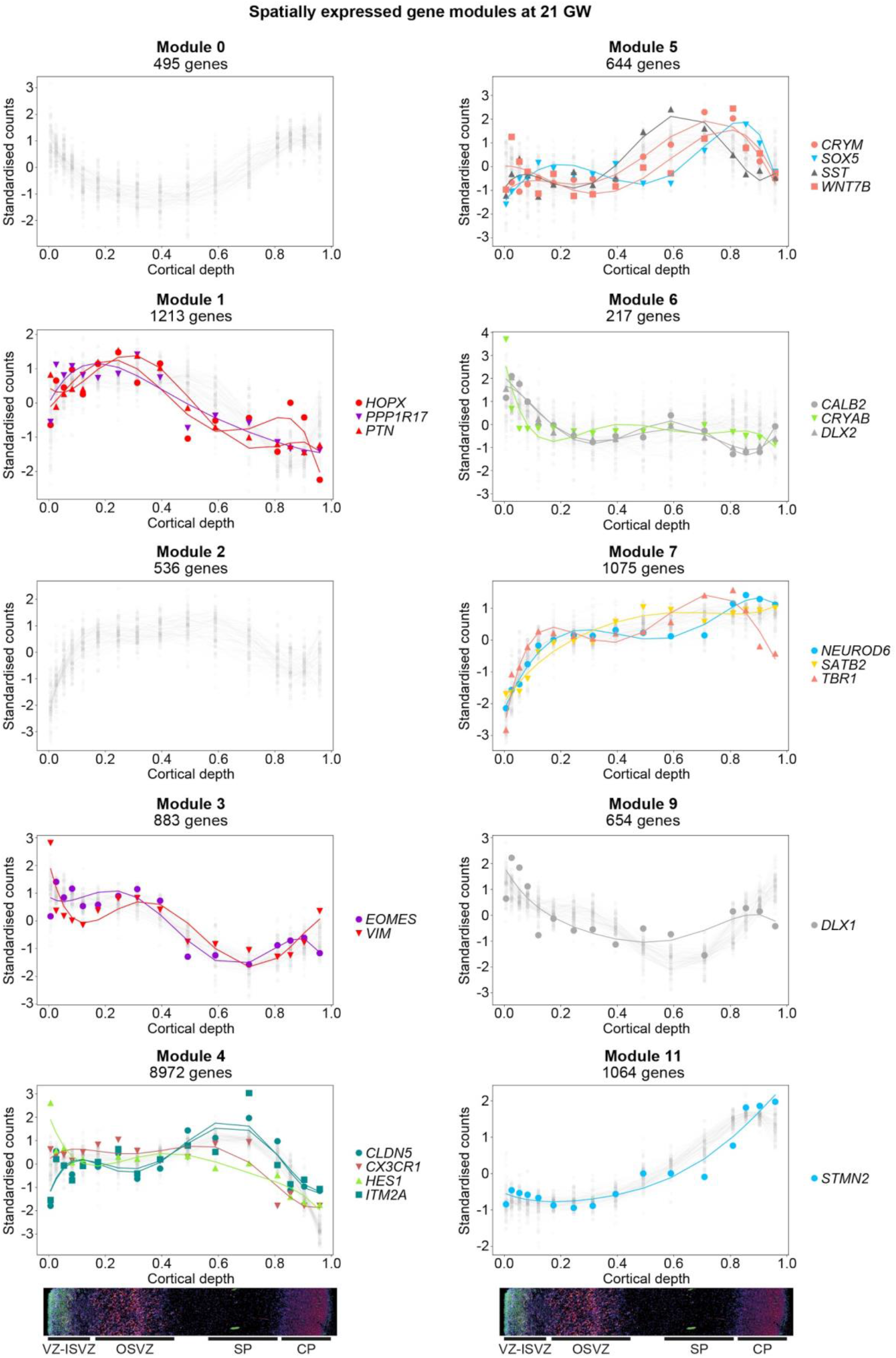
Classification of genes in the 21 GW developing cortex into spatial expression modules by SpatialDE. These were classified into 12 modules, many of which bear resemblance to specific major cell types. Two modules (8 and 10) were removed, because one contained no genes and one only contained genes below the limit of detection. See also Supp. Table 2.

**Supp. Figure 5:**
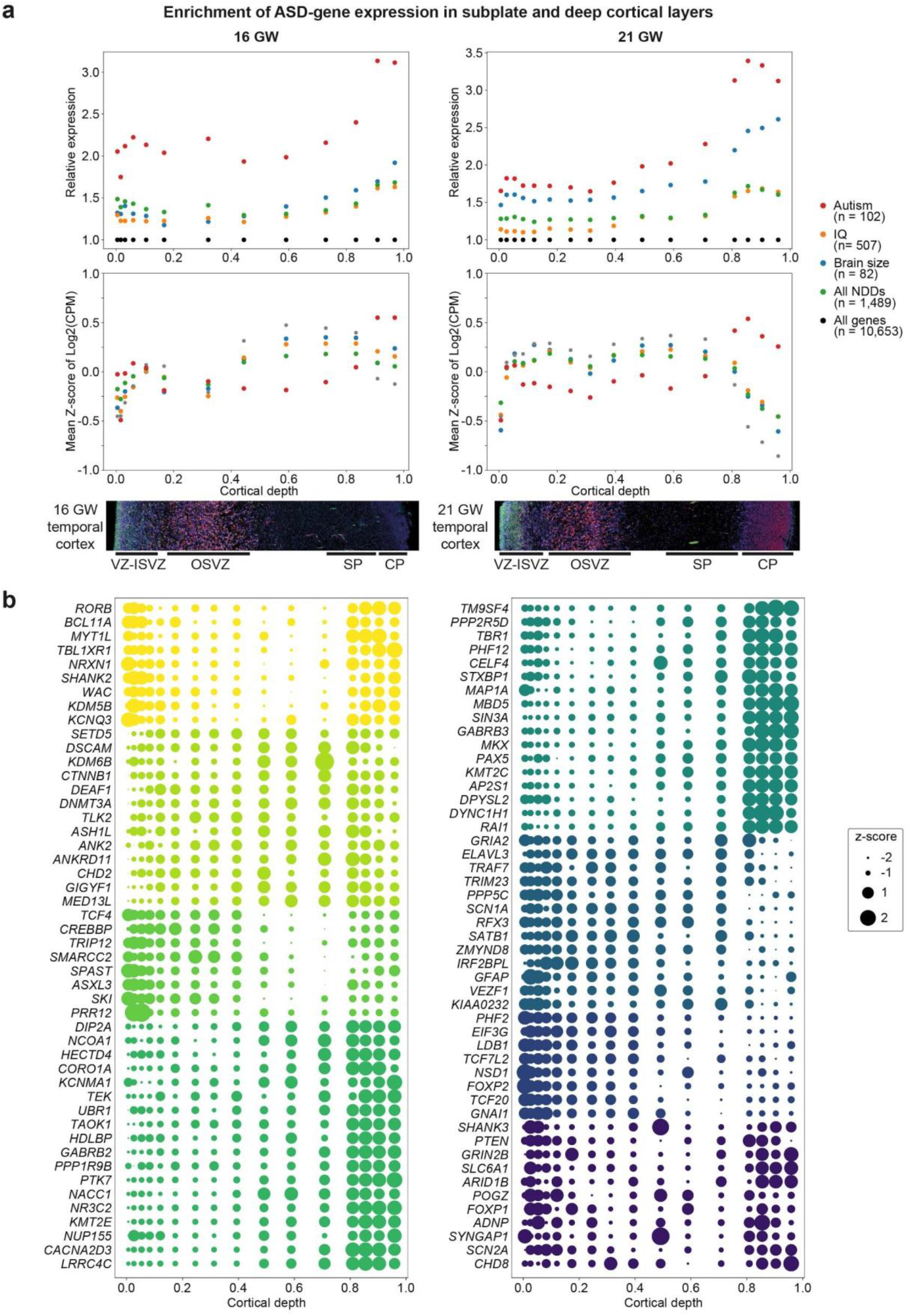
Spatial expression of ASD genes measured by WTA. **a**, Autism-related genes are spatially enriched in the subplate-deep cortical layers at both 16 GW (left) and 21 GW (right), as shown by both relative expression (top) and mean z-score (bottom). **b**, Spatially resolved expression of individual ASD genes at 21 GW, depicted as z-scores relative to all genes. Colours indicate clusters obtained from average linkage hierarchical clustering on the gene-gene correlation matrix.

**Supp. Figure 6:**
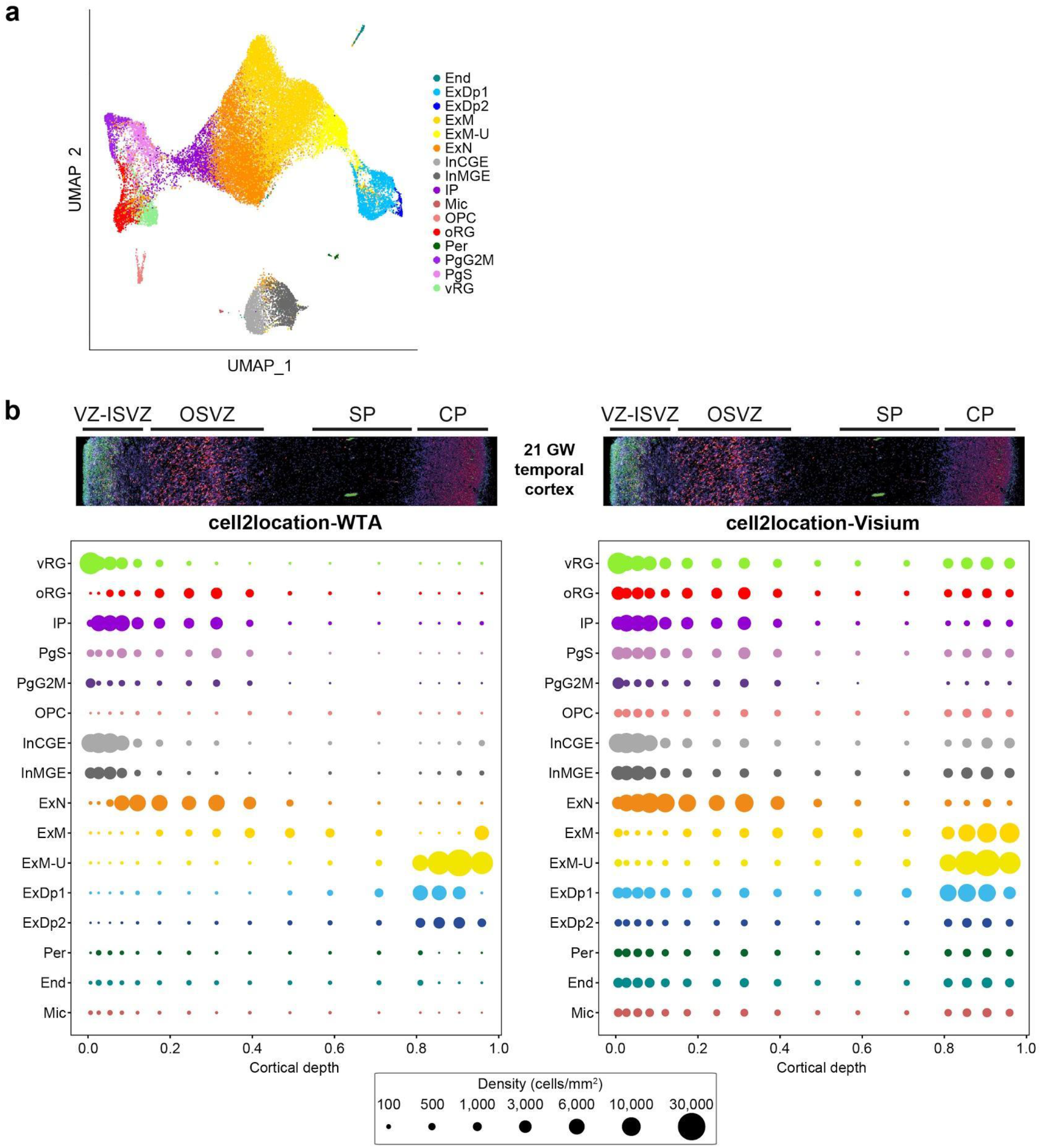
Integration of WTA with a scRNA-seq reference dataset using cell2location-WTA. **a**, UMAP plot of the scRNA-seq reference dataset used for decomposition of ROIs/AOIs, including all broad cell types of the developing neocortex at 17-18 GW^16^. **b**, Comparison of spatial mapping of broad cell types at 21 GW using cell2location, designed for 10X Visium data, and the newly adapted cell2location-WTA, which incorporates background correction by CountCorrect. cell2location-WTA provides a far more accurate mapping of broad cell types in WTA data than the cell2location-Visium model, such as the specific enrichment of progenitors (vRG, oRG, IP) in the germinal zones and maturing neurons (ExM-U) in the cortical plate.

**Supp. Figure 7:**
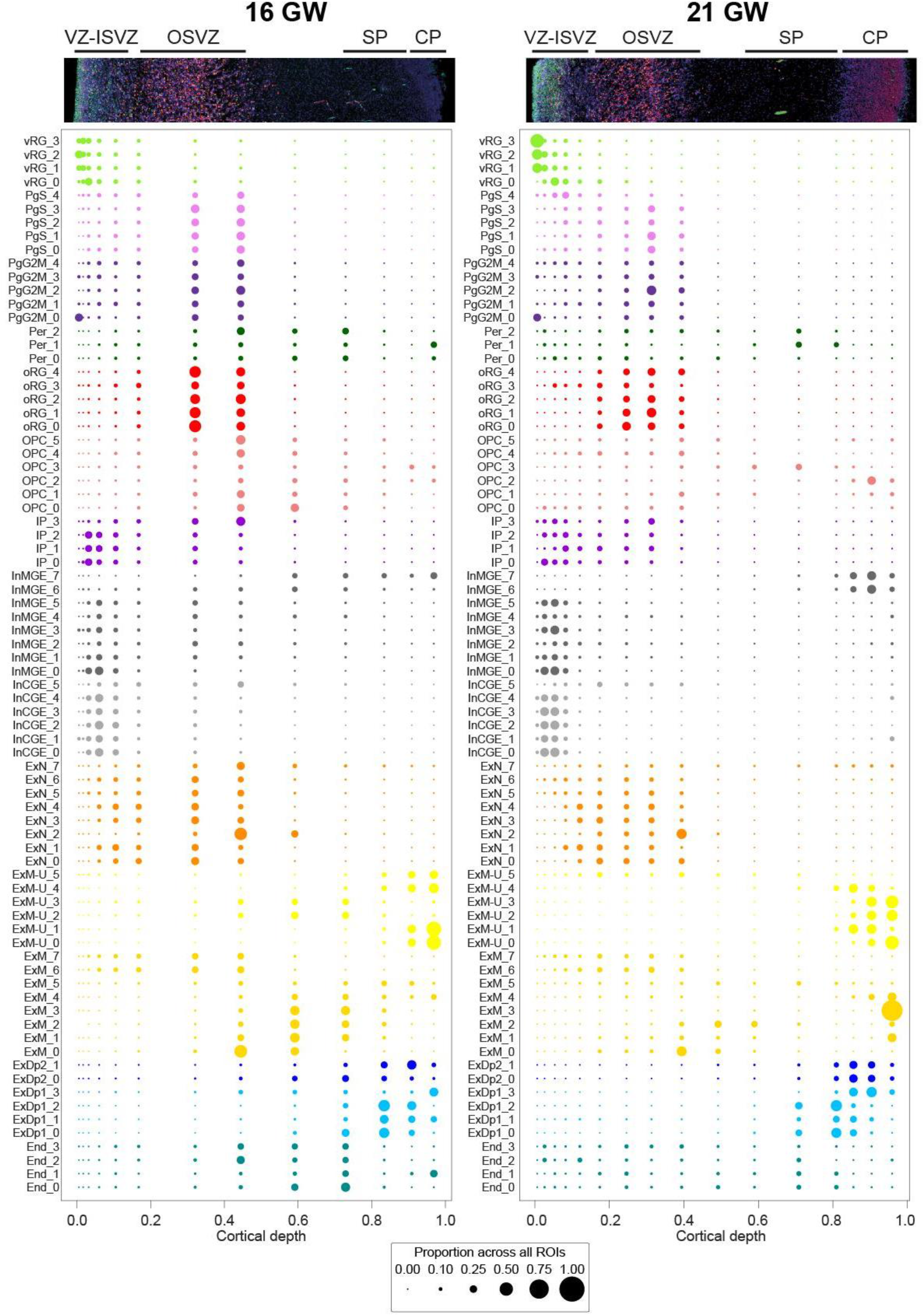
Spatial mapping of fine cell subtypes. Mapping by cell2location-WTA of all 78 cell subtypes in the scRNA-seq reference dataset^16^ at 16 GW (left) and 21 GW (right).

**Supp. Figure 8:**
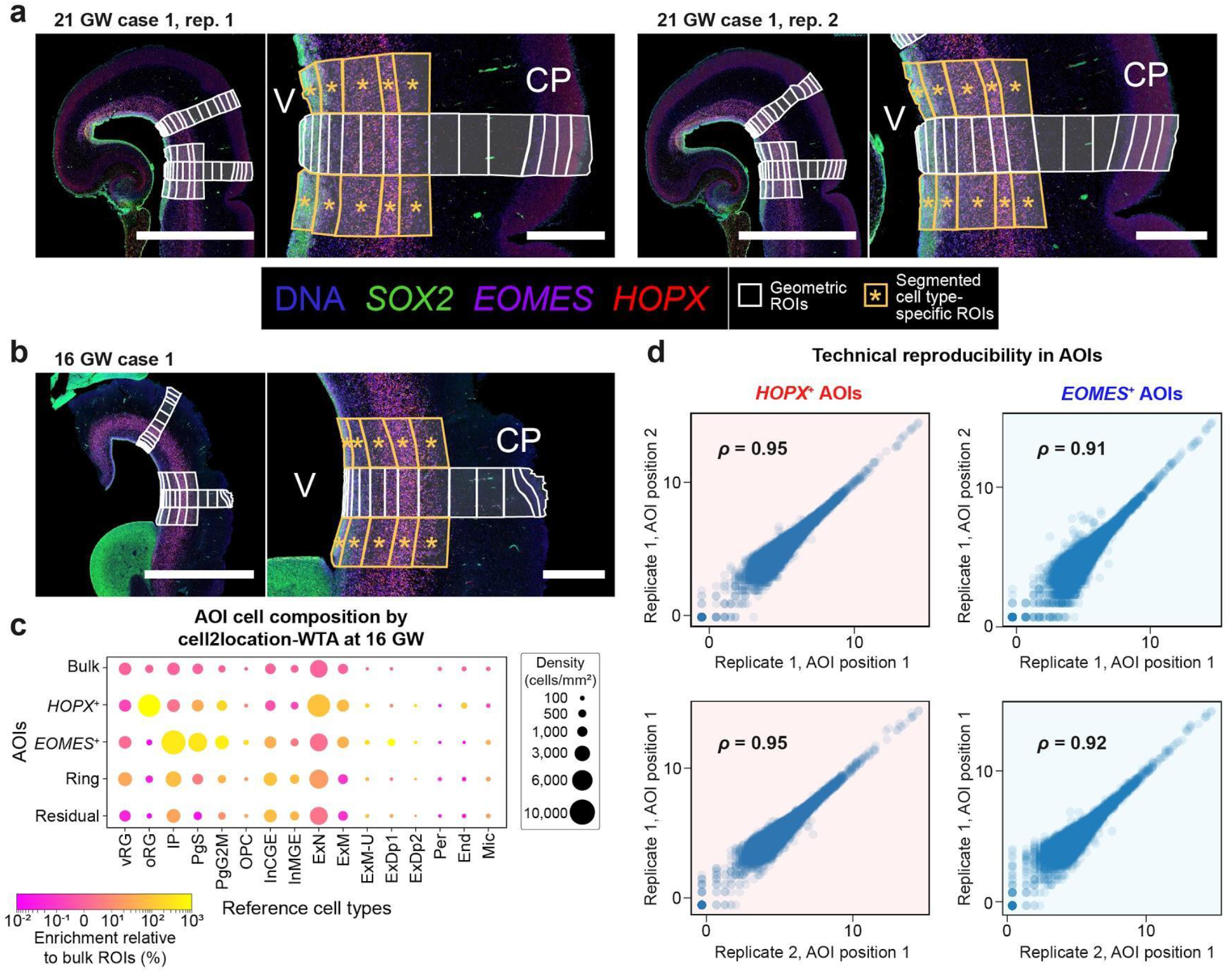
AOI segmentation across all samples. **a-b**, GeoMX custom image segmentation strategy shown for all three slides featuring segmented ROIs: two technical replicates (adjacent sections) of 21 GW case 1 (**a**); and 16 GW case 1 (**b**). Scale bars, left of each panel 5 mm, right 1 mm. **c**, Analysis of AOI cell composition at 16 GW by cell2location-WTA indicates strong enrichment of targeted cell populations: oRGs in HOPX^+^ AOIs; and IPs, PgS and PgG2M in EOMES^+^ AOIs. Shown is the mean decomposition of ten segmented ROIs, relative to neighbouring bulk ROI data. The four most superficial HOPX^+^ AOIs were excluded due to the lack of HOPX^+^ oRGs in the VZ. **d**, Technical reproducibility within AOIs, comparing transcriptome-wide expression between HOPX^+^ AOIs (left) and between EOMES^+^ AOIs (right). Top, comparison of two AOIs from the same slide; bottom, comparison of AOIs in the same position on adjacent slides. All comparisons used AOIs from the densest part of the OSVZ.

**Supp. Figure 9:**
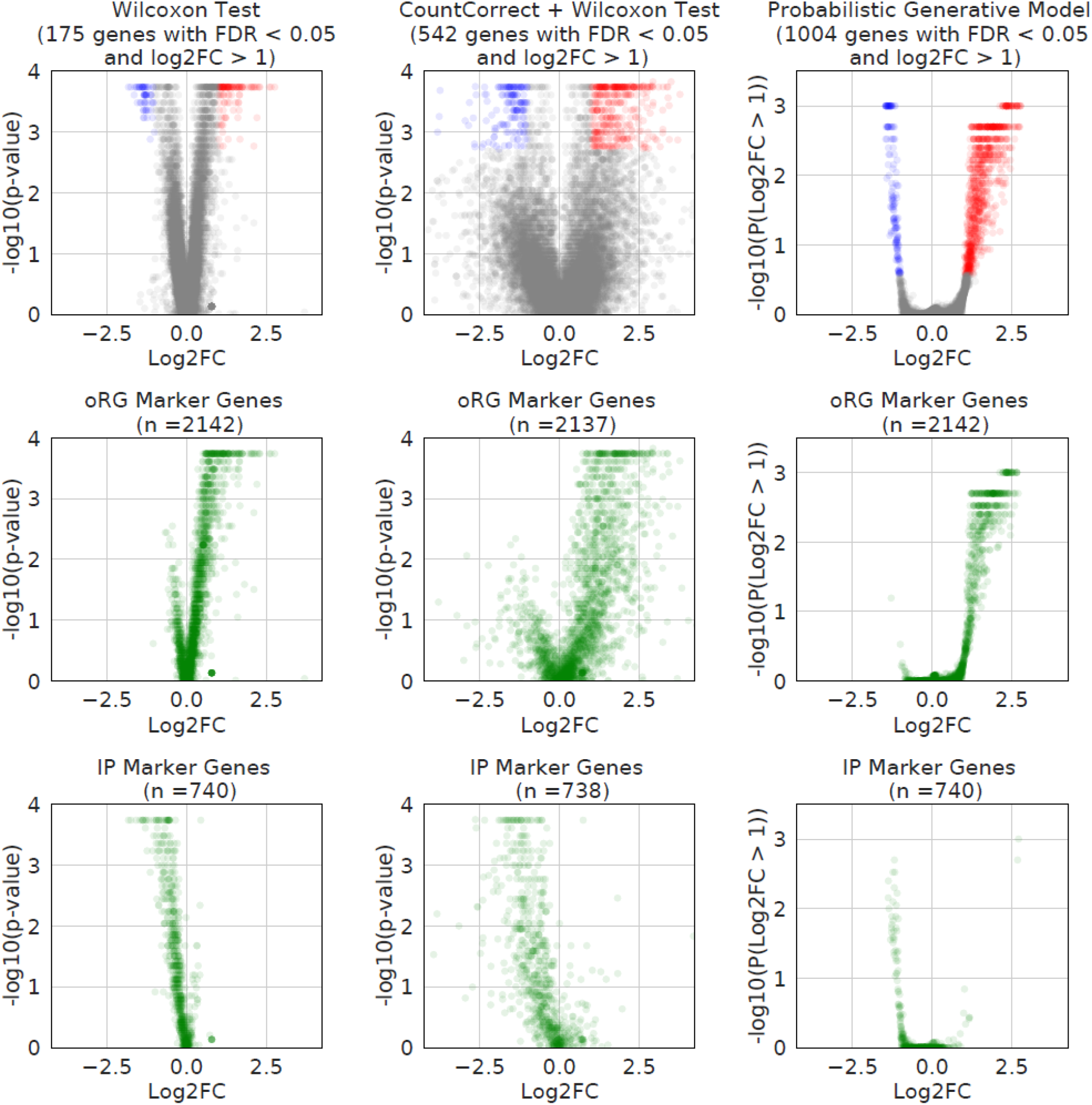
Comparison of differential expression methods on cell type specific WTA data. (Top row) volcano plots of differential expression results using the nonparametric Wilcoxon Test on raw counts (left), the nonparametric Wilcoxon Test on corrected counts (middle), or our probabilistic generative model (right). (Middle and bottom rows) placement of oRG and IP markers, respectively, extracted from scRNA-seq^16^ onto the above plots. The Wilcoxon Rank Sum Test on corrected counts was selected over the probabilistic generative model (PGM), due to the strong dependence of the latter on parameter choices. The PGM results in this figure were produced with the overdispersion parameter set to 0.001, which increases the number of differentially expressed genes over the default choices, as further discussed in the Supp. Methods.

**Supp. Figure 10:**
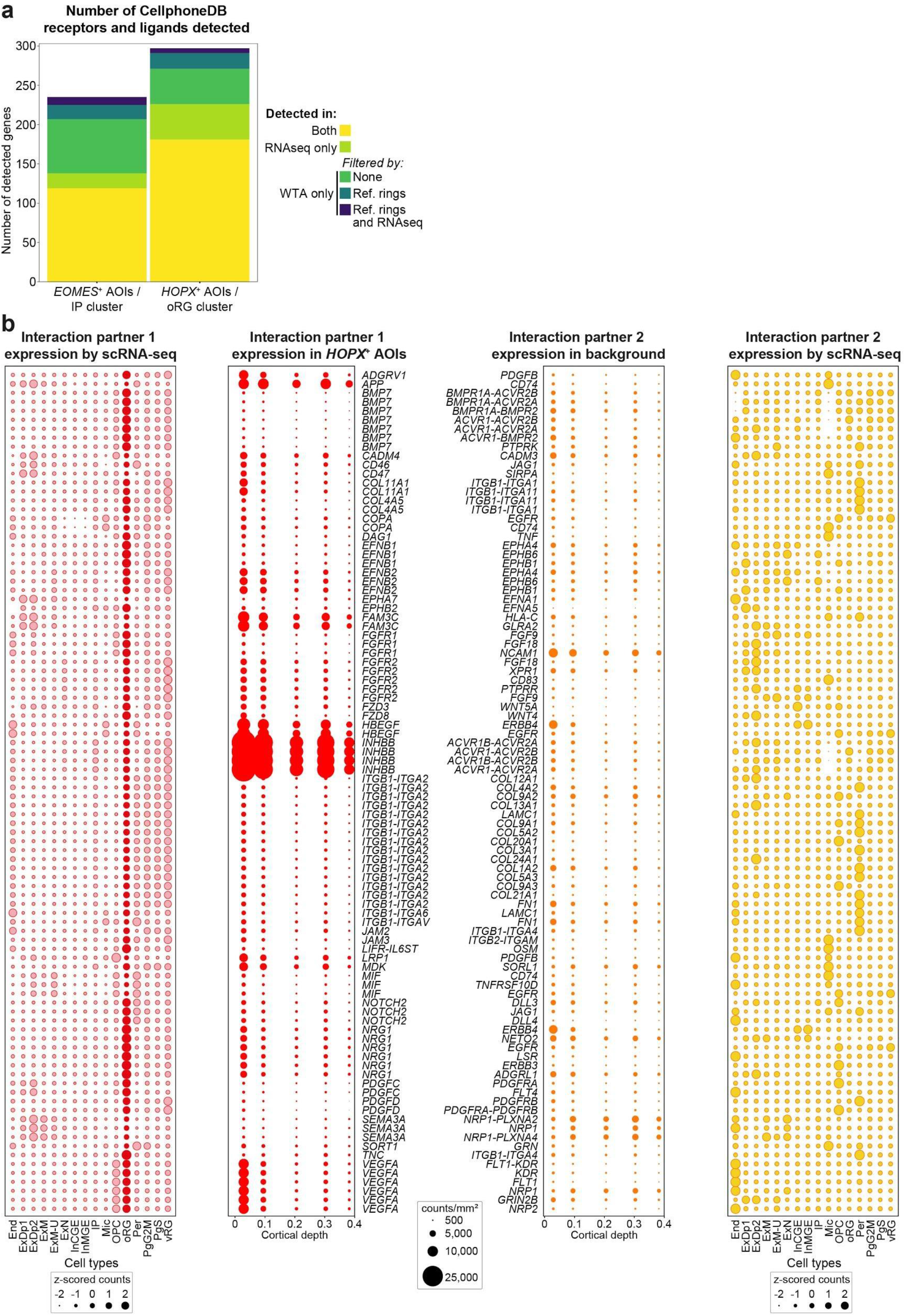
Cell interaction analysis using cell type specific WTA. **a**, Stacked bar plot showing the detection of CellPhoneDB receptors or ligands in WTA and the scRNA-seq reference. All CellPhoneDB genes detected in one or both of the RNAseq reference and the Nanostring WTA probe panel are considered. For RNAseq a receptor/ligand was judged to be detected if it had at least 0.02 mean counts in the IP or oRG cluster. For WTA, detection was defined as exceeding the limit of detection in at least 2 of 20 AOIs, defined as the mean plus 2.5 standard deviations of negative probe counts calculated on a log2 scale. Unique WTA genes were filtered in two ways to exclude bleedthrough from other cell types. Firstly, requiring the gene to be expressed more highly in HOPX^+^/EOMES^+^ AOIs than in reference rings in more than 50% of ROIs; second, additionally requiring the gene to be undetected in all cell types in RNAseq data. Using the most stringent filtering for the WTA ligands/receptors, the Jaccard index would be 0.70 for EOMES^+^ AOIs/IP cluster and 0.66 for HOPX^+^ AOIs/oRG cluster. **b**, oRG specific interactions in scRNA-seq and WTA. From the initial set of 124 interactions enriched in oRGs (cellPhoneDB p-value < 0.05) 113 were detected in WTA. Out of 11 missing interactions, six included receptors or ligands that were not part of the WTA probe panel leaving only five interactions that included genes that were profiled, but not detected in WTA (Supp. Table 3). The plot shows the 94 interactions between oRGs and other cell types (i.e. oRG-oRG interactions not shown). Central dotplots show WTA expression derived from 21 GW biological case 1, technical replicate 1 HOPX^+^ AOIs and background (residuals and EOMES^+^ AOIs). Outer dotplots on left and right show z-scored expression of interaction partners 1 and 2, respectively, in scRNA-seq data^16^.

**Supp. Figure 11:**
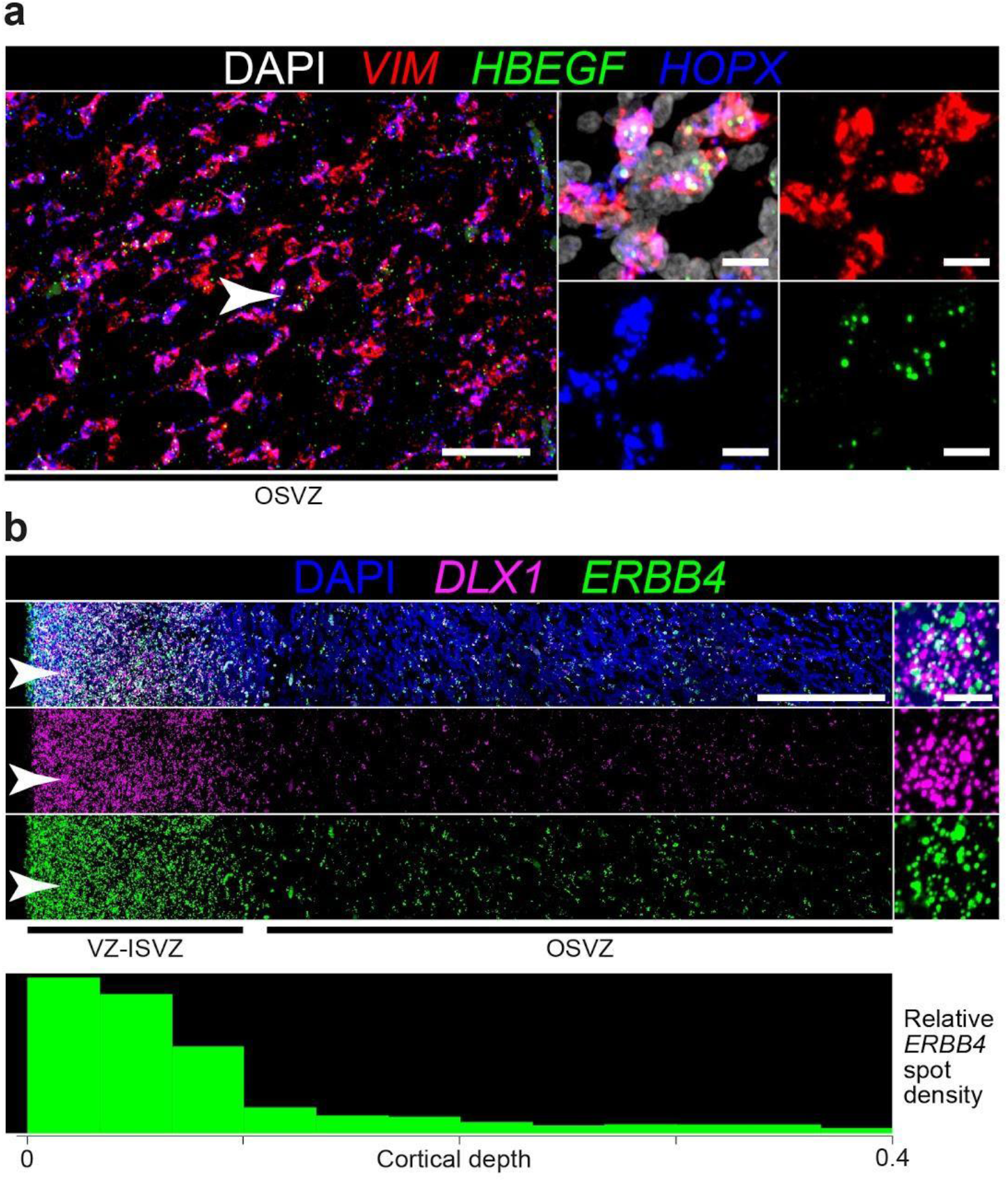
Spatial validation of the HBEGF-ERBB4 interaction. **a**, Validation by RNAscope smFISH of HBEGF expression in HOPX^+^ VIM^+^ oRGs in the OSVZ at 21 GW. Scale bars, left 200 μm, right 20 μm. **b**, Validation of spatial expression of ERBB4 in superficial germinal zones by spot quantification of RNAscope smFISH. ERBB4 is co-expressed with the developing interneuron marker DLX1. Scale bars, left 500 μm, right 10 μm.

## Supplementary Methods

## 1 Nanostring WTA Generative Model

Here we present our generative model of both expression signal and measurement noise for the Nanostring WTA platform.

We assume that measurement noise is composed of “sampling noise”, arising from incomplete detection of all RNA molecules in a sample, and “background noise” arising from non-specific binding of each probe to non-target transcripts. Sampling noise can be accurately modelled with a Poisson distribution for UMI count data if the number of targeted genes is large and the detection probability is low [12]. Both assumptions are fulfilled in Nanostring WTA data. Additional variability, for example arising from biological heterogeneity not accounted for in the model, can be modelled with a Negative Binomial distribution [12]. Unlike the Poisson distribution that has a variance equal to the mean, the Negative Binomial distribution can include an additional gene-specific variance, *v*_*g*_.

For an ROI or AOI with a single cell, WTA UMI counts, *X*_*rg*_, would thus arise from a simple Negative Binomial distribution:

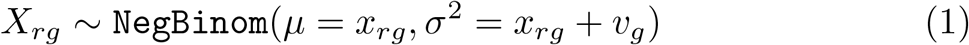

where the first subscript *r* denotes ROIs/AOIs and the second subscript *g* denotes genes.

For an ROI with *m*_*r*_ cells/nuclei, the mean changes by a factor of *m*_*r*_ and the variance by a factor 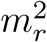:

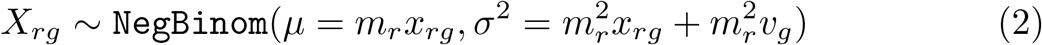

We then decompose the mean into two parts given by the sum of specific, *A*_*rg*_, and non-specific probe binding, *B*_*rg*_:

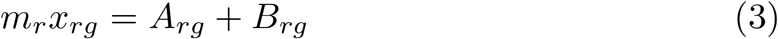

Since we typically will not have enough samples to estimate the extra variance *v*_*g*_ for each gene accurately, we pool information across all genes with similar expression. We achieve this by drawing all *v*_*g*_ from a common prior Gamma distribution. This distribution has a variance and a mean that scales linearly with the average expected real counts of each gene, per nuclei, which we call *r*_*g*_.

First, *r*_*g*_ is given by:

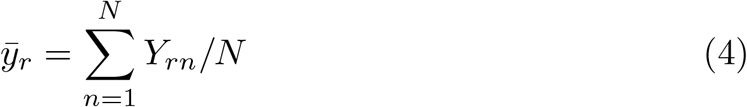

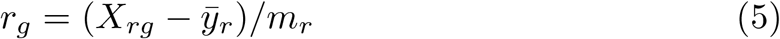

where *Y*_*rn*_ are the negative probe counts.

Next *v*_*g*_ is defined as:

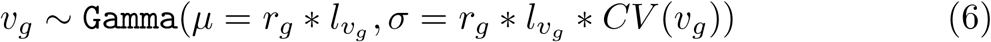

with *CV* (*v*_*g*_) = 0.01 and the following linear scaling parameters 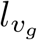:

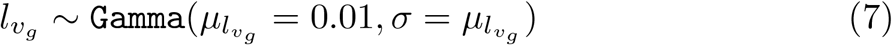

We aim to avoid biologically unlikely models in which all gene expression is explained by extra noise, i.e. high and different overdispersion values for each gene. This is why we choose small prior values of 0.01 for *CV* (*v*_*g*_) and 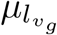.

We now describe our model for non-specific binding. The counts from *N* =138 negative probes can provide a first estimate of the background noise in each ROI/AOI. We note that negative probe counts increase linearly with the total number of RNA counts in a sample (figure 2, 3), with a different linear slope for each negative probe (figure 4). With *l*_*r*_ defined as the total number of counts in an ROI, an appropriate model for the negative probe counts *Y*_*rn*_ would thus be:

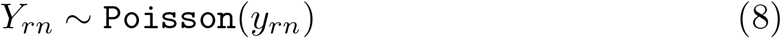

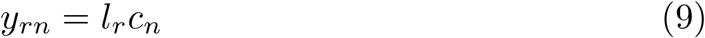

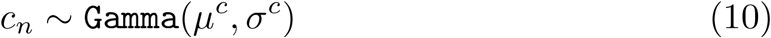

where the subscript *n* denotes negative probe counts.

The empirical mean of negative probe counts normalized by total counts is given by:

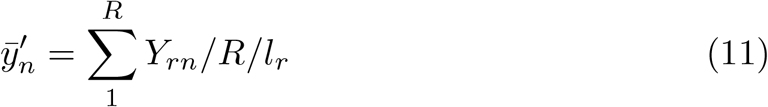

where *R* is the total number of ROIs.

**Figure 1:**
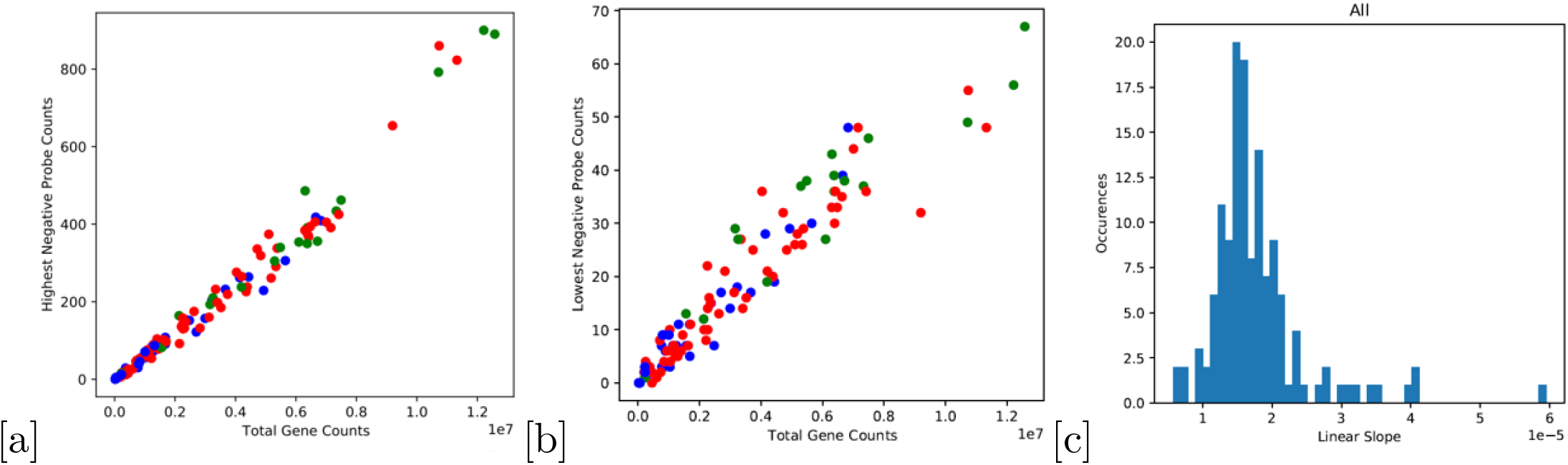
[a] An example of counts for one negative probe across different ROIs/AOIs (different slides indicated by different colours). This example probe is the one with highest counts overall. A strong linear relationship with total counts in each ROI/AOI is evident. The slope is 590 counts per 10^7^ total counts. [b] An example of counts for one negative probe across different ROIs/AOIs (different slides indicated by different colours). This example probe is the one with lowest counts overall. A linear relationship with total counts in each ROI/AOI is evident. The slope is 45 counts per 10^7^ total counts. [c] The distribution of linear slopes for all negative probes.

We use it to define our expectations for *µ*^*c*^ and *σ*^*c*^:

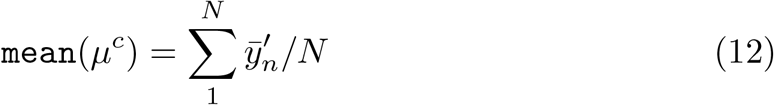

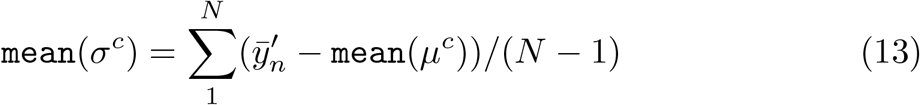

We use these values to initialize the prior distribution parameters for *µ*^*c*^ and *σ*^*c*^. Since we cannot be sure that the negative probe counts are an accurate representation of non-specific binding that also holds true for gene probes, we add an additional uncertainty corresponding to a coefficient of variation of 0.25:

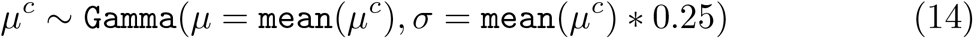

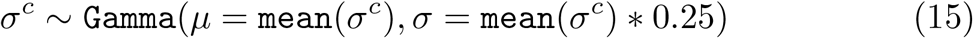

We then assume that non-specific binding of gene probes results from the same generative process as binding of negative probes, and can be modelled as:

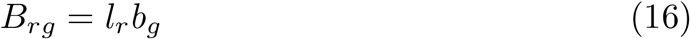

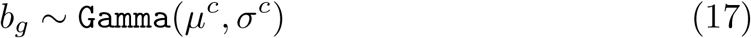

For the specific-binding component *A*_*rg*_ we build on existing methods in the field of single-cell transcriptomics (e.g. [5, 6, 7,9, 11]), which assume that gene expression can be accurately described in a low-dimensional space. Biologically this assumption rests on the existence of discrete cell types and transcriptional modules of highly correlated genes. We chose non-negative matrix factorization with 15 factors, *f*, as our dimensionality reduction method, so that *A*_*rg*_ is given by:

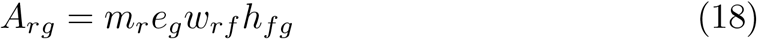

where *w*_*rf*_ are the weights for the 15 factors in each ROI/AOI, *h*_*fg*_ are gene loading values for each factor and *e*_*g*_ is a gene scaling factor that is useful for interpreting the model, as we show below. The components have hierarchical priors given by:

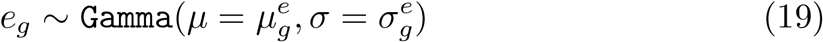

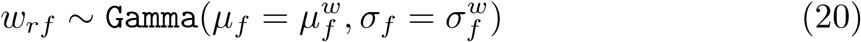

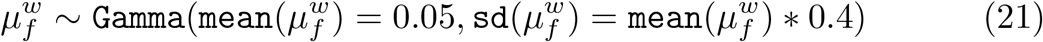

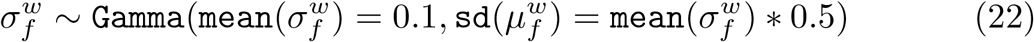

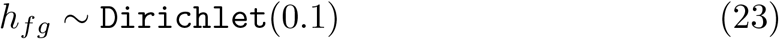

Our prior parameter choices for the *w*_*rf*_ and *h*_*fg*_ components, result in sparse matrices, i.e. most entries of *w*_*rf*_ and *h*_*fg*_ are expected to be 0. This choice of prior parameters gives higher weight to simple and biologically meaningful solutions with few active components. Moreover, the identified components are each active in only a few ROIs and contain only a fraction of the total number of genes in the probe panel. In turn it avoids complex and biologically unlikely solutions, where gene expression is explained with a large number of ubiquitously expressed, whole-transcriptome components. Importantly, the hierarchical prior distribution of *w*_*rf*_ should result in automatic relevance detection as in [10], so that if a given dataset is fit much better with a large number of factors, the prior mean for *w*_*rf*_ will converge on a larger value and most or all factors can be “switched on” (i.e. obtain non-zero weights).

Given the Dirichlet distribution for the gene loadings the following relation holds for all genes:

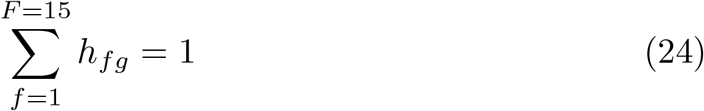

Consequently, differences in total gene expression can be modelled with the *e*_*g*_ parameter instead of *h*_*fg*_. The prior mean for *e*_*g*_ should be set so that the expected mean of each gene in the *A*_*rg*_ matrix corresponds to the sample mean in the data *X*_*rg*_ minus the expected background *B*_*rg*_, i.e. we require:

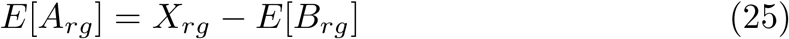

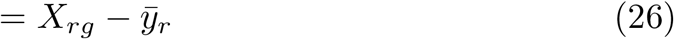

We do this by first considering the expected mean of the *A*_*rg*_ matrix:

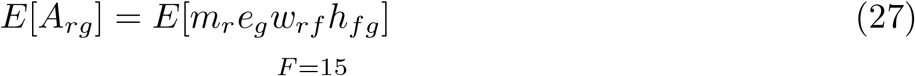

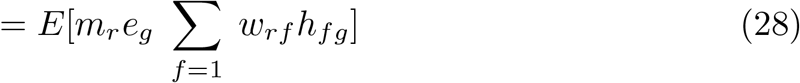

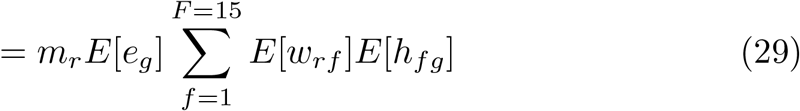

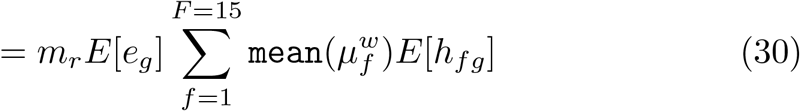

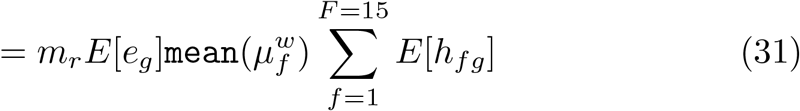

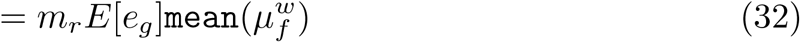

where we used equation 24 to move from 29 to 30. Now equating 26 and 32 and rearranging we obtain the appropriate mean for the prior distribution of *e*_*g*_:

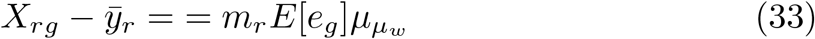

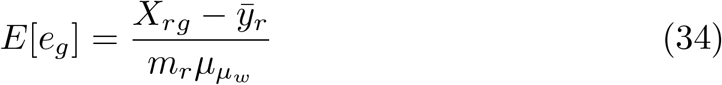

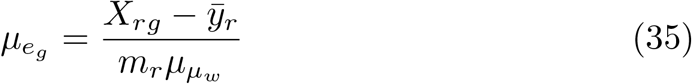

We set the CV for the *e*_*g*_ prior distribution to 0.25:

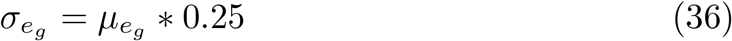

We show later that this CV parameter has a large effect on the model. Together with the background CV in equation (14) it determines to what extend we rely on the negative probe counts to estimate the magnitude of non-specific binding. A CV of 0 would not allow the model to converge to any other solution that then trivial solution of estimating a real expression and background based on the mean negative probe counts. A value of 1 or larger would give almost no weight to the negative probe counts and would estimate the magnitude of specific and non-specific binding based solely on the best fitting non-negative matrix factorization. We find that a value of 0.25 falls between those two extremes (see section 7 on robustness).

## 2 Prior predictive check

We confirmed numerically that the prior values we have specified for *e*_*g*_ result in a prior distribution that corresponds well to the values for *X*_*rg*_ found in the real data. Figure 2 shows the prior sample mean of *X*_*rg*_ for each gene on the x-axis and the real mean counts across ROIs on the y-axis. Since the two values match up for all genes our prior specifications of *X*_*rg*_, we conclude that the specific (*A*_*rg*_) and non-specific binding binding component in our model (*B*_*rg*_) are consistent with the data.

**Figure 2:**
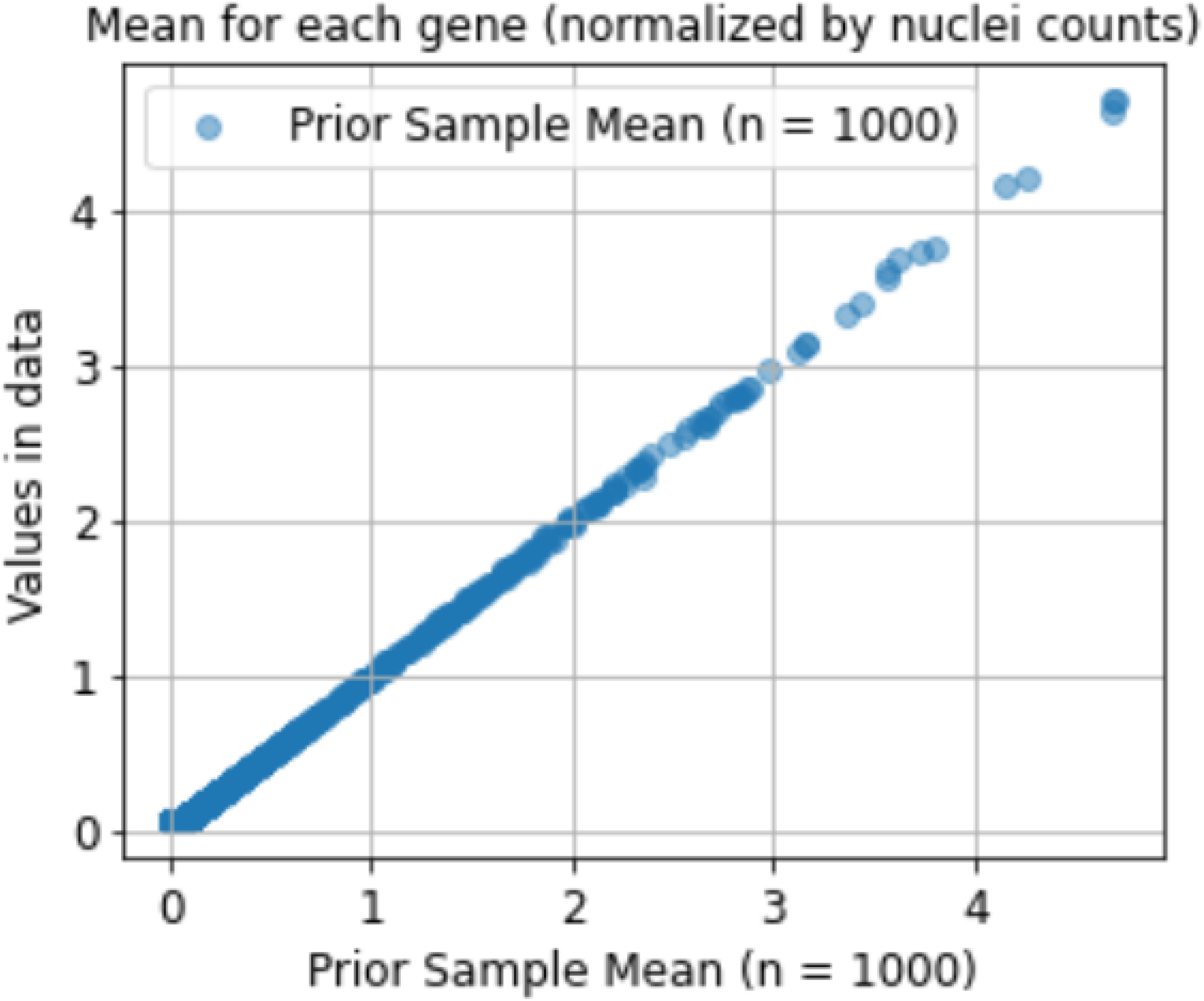
Using 19 pcws biological and technical replicate 1, the figure shows the mean counts of each gene in the data on the y-axis, plotted against its prior expected expression on the x-axis. Both values are normalized for nuclei counts. By inspection values match up for each gene, so our prior expectations for the specific (*A*_*rg*_) and non-specific binding binding component in our model (*B*_*rg*_) are consistent with the data.

## 3 Inference

We implemented the model described in section 1 in the pymc3 probabilistic programming language [8]. We then used mean-field Automatic Differentiation Variational Inference (ADVI) [4] implemented in the pymc3 framework to obtain the approximate posterior distribution of the parameters in our model. Like other variational inference algorithms, ADVI reframes the inference procedure as an optimization problem by defining a family of approximating distributions for the latent variables and minimizing their KL-divergence to the posterior by maximizing the expectation lower bound (ELBO). Briefly, ADVI firstly employs effective transformations of the latent parameters, so that their distributions can be accurately modeled by factorized, standardized Gaussian distribution in the transformed space. Secondly, it uses MCMC to obtain expectations of the ELBO at each step, which can then be used in stochastic optimization methods. Pymc3 only requires the specification of the model, an optimizer, learning rate, and iteration number to perform ADVI. In our case, we chose the ADAM optimizer, a learning rate of 0.001 and 300,000 iterations. We ran the optimization on each of the 4 data batches (corresponding to 4 experiment slides) separately. We visually checked for convergence of the ADVI optimization by plotting the objective function (ELBO) over all 300,000 iterations, as well as the last 10,000 iterations, for which we show an example in figures 3 and 4 for the 19 pcw biological case 1, technical replicate 1.

**Figure 3:**
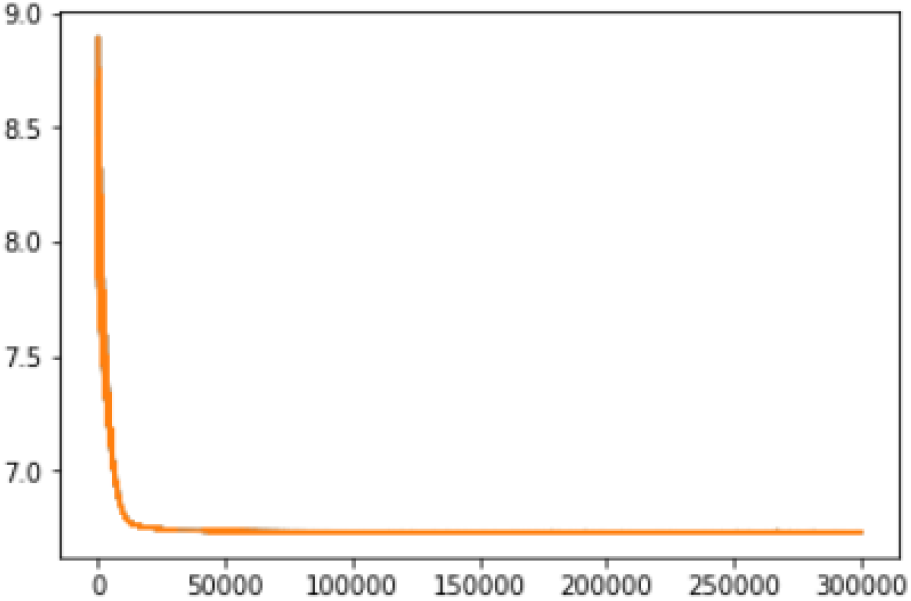
Example of optimization history across all 300,000 iterations. 19 pcw biological case 1, technical replicate1.

**Figure 4:**
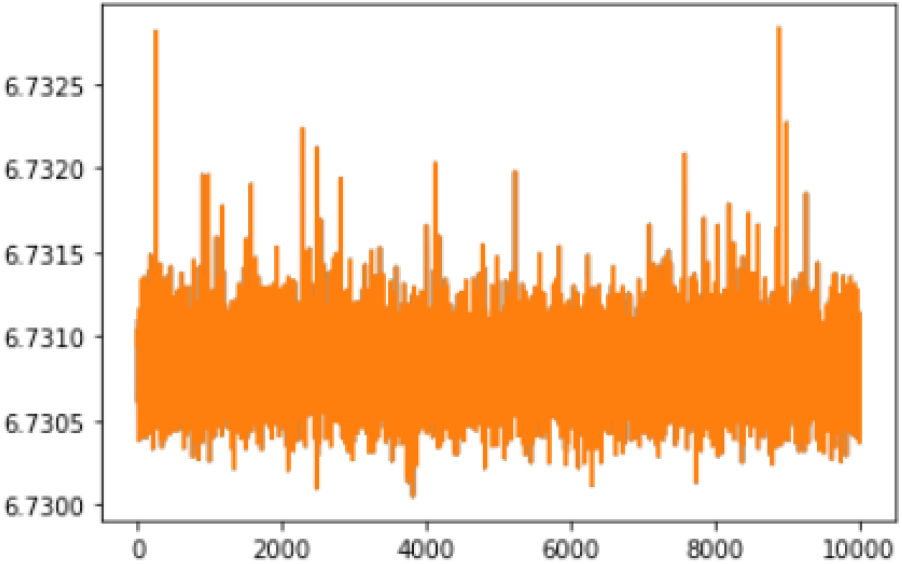
Example of optimization history across the last 10,000 iterations. 19 pcw biological case 1, technical replicate 1

## 4 Posterior approximation

For all downstream applications we use 1,000 samples from the ADVI distribution to approximate each parameter’s posterior distribution. We then obtain the mean and variance of each parameter from the sample mean and variance across all 1,000 samples.

## 5 Denoising

There are two different approaches to obtain a denoised dataset, which we call 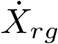 and 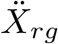.

We can either subtract expected background counts from the count matrix:

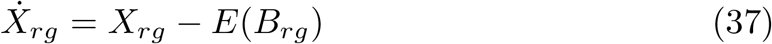

Or define corrected counts as the expected low-rank representation of real counts *A*_*rg*_:

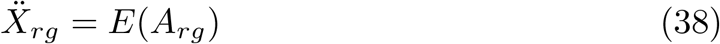

The latter approach can remove sampling noise on top of background counts, but may also remove some biological variation.

By default we use the first approach in our CountCorrect method, round all values to integer values and clip values at a minimum of 0.

## 6 Differential expression analysis

The ADVI method returns an approximation for the full posterior distribution so that for downstream analysis tasks, such as differential expression analysis, we could in principle not only use the mean estimates -which are returned by our CountCorrect method -but also the variance to construct a more accurate differential expression method. Previous work in single-cell transcriptomics analysis [1] followed such a strategy that combined a probabilistic generative model and variational inference to perform differential expression analysis. We also constructed such a method to work on the full ADVI posterior distribution, but found it to be less robust than simply using the mean estimates returned by CountCorrect in combination with a standard Wilcoxon test (see next section on robustness). For completeness we still explain our differential expression method below.

Our goal is to determine the probability, *p*_*g*_, that the log2-fold change in expression, 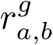, between ROI *a* and *b* for a gene *g* is greater than a certain threshold *δ*, based on the CPM normalized 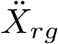 matrix:

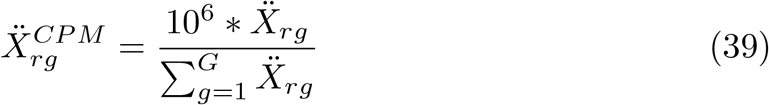

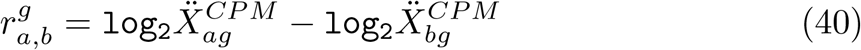

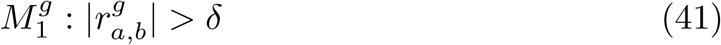

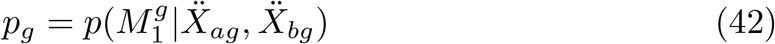

If *a* and *b* are not single ROIs, but two groups of ROIs, we consider the difference in mean gene expression across the two groups instead. We calculate the probability distribution for 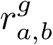 numerically based on the 1,000 samples from the 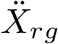 posterior distribution. Similarly, we then calculate a probability that the log_2_-fold change is greater than a certain threshold by summing the probability density above this threshold (i.e. the number of samples above this threshold, divided by the total number of samples). For calculating the expected false-discovery rate (FDR) we use the formula from [2]:

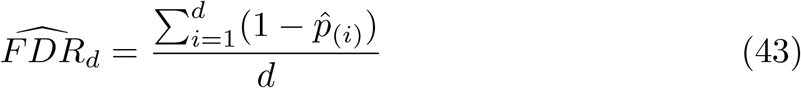

where 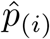 are the probabilities of differential expression for all genes in decreasing order and *d* is the rank of the probability for which we calculate the corresponding FDR.

In principle, both 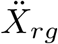 and 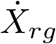 could be used for this method. Notably, we use the low-rank representation of real counts 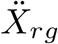 from equation 38 for this purpose and not the subtraction method, 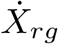 from equation 37. This is because 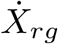 would only model uncertainty in the background and thus underestimate the uncertainty in the real counts.

## 7 Robustness

We compared three methods of differential expression analysis with respect to their robustness to parameter changes in the generative model and chose the CountCorrect+Wilcoxon test method presented in the main text, because it emerged as the most robust. The differential expression analysis was performed between HOPX+ and EOMES+ AOIs from biological case 1, technical replicate 1 as described in the main text. The three methods we considered were:

1. Wilcoxon-CC1: Using the Wilcoxon test on CountCorrect cpm-normalized counts.
2. Wilcoxon-CC2: Using the Wilcoxon test on CountCorrect cpm-normalized counts, but with CountCorrect using equation 38 instead of the 37 to return denoised counts.
3. PGM-CC2: using the probabilistic generative model approach described in the previous section.

The four parameters we varied were:

1. *n*-factors: The number of factors used in non-negative matrix factorization, as defined in equation. Be default this value is set to 15. We swept through values 5,10,15,20 and 50.
2. 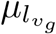: The mean of the prior distribution for the 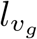 overdispersion parameter in equation. Since the coefficient of variation of the prior distribution is fixed at 1 this parameter also affects the variance of the prior distribution. By default this value is set to 0.01. We swept through values 0.001 to 0.25.
3. CV-background: The coefficient of variation of prior background mean *µ*_*c*_ and variance *σ*_*c*_, as defined in equations and. Be default this value is set to 1. We swept through values 0.01 to 25.
4. CV-expression: The coefficient of variation for the prior distribution of the *e*_*g*_ parameter, as defined in equation. The default value is 1. We swept through values of 0.01 to 1.

The results of this analysis are shown in figures 7 to 10. Only one parameter, “CV-expression”, resulted in large changes in the number of differentially expressed genes detected with the Wilcoxon-CC1 method as shown in figure **??**. In particular, if the prior uncertainty in the real expression is increased, the number of differentially expressed genes increases. We found that this is a direct result of the model estimating a higher average background level than suggested by the negative probes since the prior expected real expression is related to the background in equation 35. The “CV-expression” parameter can thus be set to higher values if the user has little confidence in the negative probe counts to represent the real magnitude of the background in the experiment. We decided to choose a moderate value of 0.25, so that overall our default settings for the CountCorrect model are as shown in section 1: n-factors = 15, 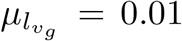, CV-background = 0.25, CV-expression = 0.25

## 8 Cell2location-WTA Model

The cell2location model[3] for cell type mapping decomposes location-level gene expression counts into a set of predefined signatures of reference cell types, taking into account both technology differences and location and gene specific background noise. In the following we explain our adaption to the cell2location model developed for spatial transcriptomics data, such as Visium and Slide-Seq V2, using the notation from the original cell2location publication. Specifically, in the standard cell2location model gene expression counts *d*_*sg*_ are assumed to follow a Negative Binomial distribution:

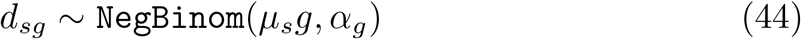

with a mean *µ*_*sg*_ given by:

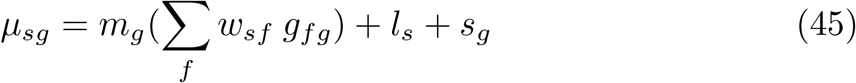

where *m*_*g*_ is a gene specific scaling parameter that adjusts for differences between technologies, *g*_*fg*_ is a matrix of reference signatures, usually obtained from single-cell RNAseq data, *w*_*sf*_ are regression weights for each reference signature and *l*_*s*_ is a “spot”-dependent (in Visium data/Slide-Seq V2) i.e. location dependent background term and *s*_*g*_ is a gene-specific background term.

To obtain the cell2location-wta method, we now only change the gene specific background term to scale with the total number of counts *t*_*s*_:

**Figure 5:**
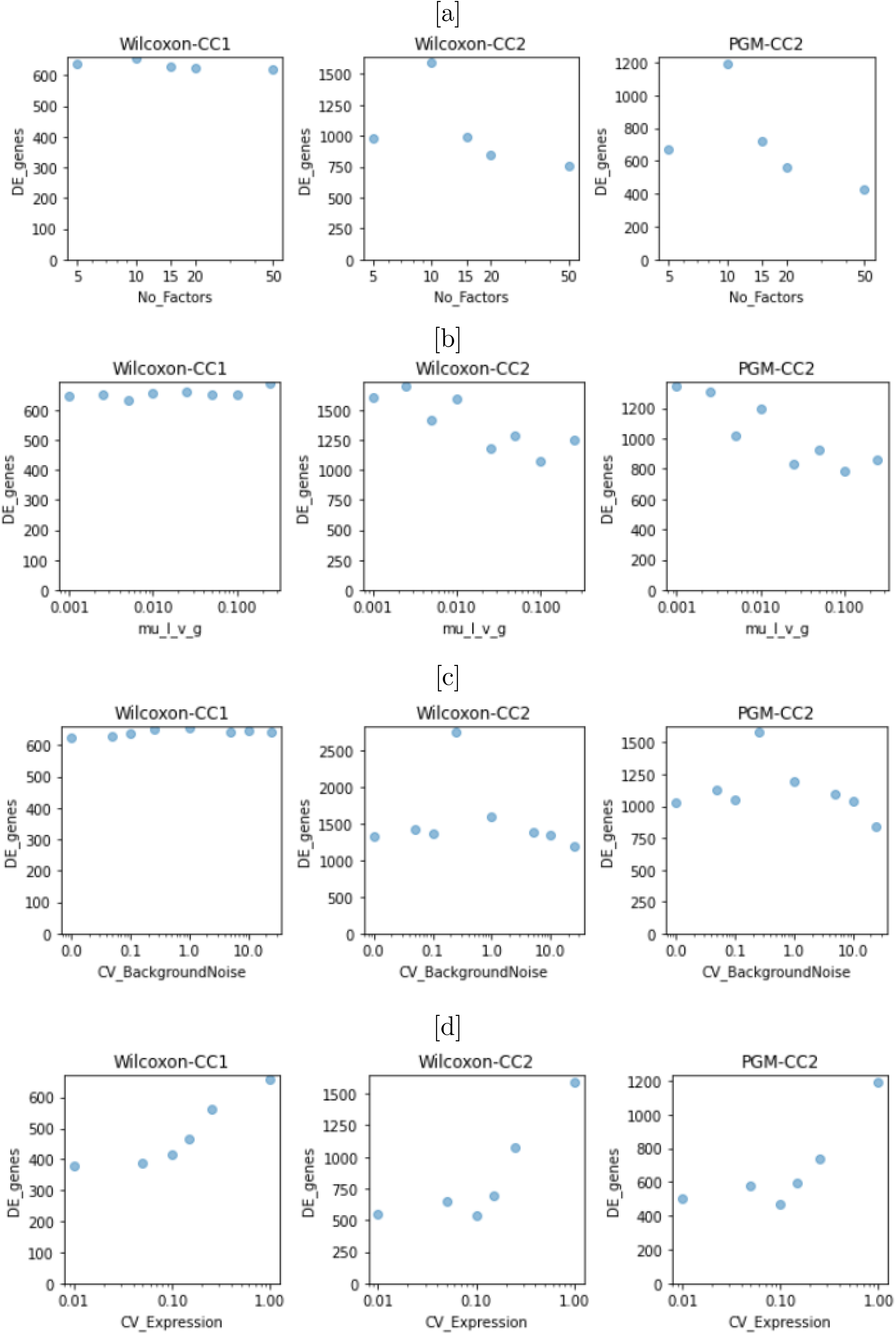
[a], [b], [c]: For our chosen differential expression method (Wilcoxon- CC1), the number of returned genes does not vary much as the number of factors, the prior uncertainty in the background noise or the overdispersion paramter is varied. [d] However, for all methods, the number of returned genes increases when we rely less on prior knowledge (negative probe counts) to determine the real expression level and more on the structure found in the data.

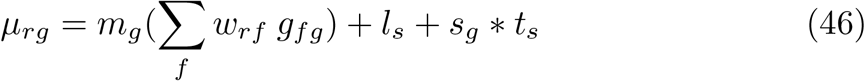

*t*_*s*_ thus corresponds to *l*_*r*_ in the CountCorrect method and the corresponding prior distribution for *s*_*g*_ is identical to the distribution for *b*_*g*_ in equation 17:

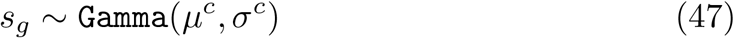

Finally, cell2location-wta, like CountCorrect, uses the negative probe counts to find *µ*^*c*^ and *σ*^*c*^ as described in equations 8 to 3.

For further details on the cell2location method we refer the reader to the cell2location publication [3].

## 9 Code Availability

The CountCorrect method is implemented as a python package and includes tutorials: https://github.com/BayraktarLab/CountCorrect The cell2location-WTA model is integrated into the cell2location package https://github.com/BayraktarLab/cell2location/blob/master/cell2location/models/LocationModelWTA.py, so that it can be used by specifying it as a non-default model choice in the main cell2location workflow as explained in a corresponding notebook that is available on the cell2location github repository as well: https://github.com/BayraktarLab/cell2location/blob/master/docs/notebooks/cell2loation_for_NanostringWTA.ipynb

## References

1. Lein, E., Borm, L. E. & Linnarsson, S. The promise of spatial transcriptomics for neuroscience in the era of molecular cell typing. Science 358, 64–69 (2017).

2. Moffitt, J. R. et al. Molecular, spatial, and functional single-cell profiling of the hypothalamic preoptic region. Science 362, (2018).

3. Qian, X. et al. Probabilistic cell typing enables fine mapping of closely related cell types in situ. Nat. Methods 17, 101–106 (2020).

4. Eng, C.-H. L. et al. Transcriptome-scale super-resolved imaging in tissues by RNA seqFISH. Nature vol. 568 235–239 (2019).

5. Ståhl, P. L. et al. Visualization and analysis of gene expression in tissue sections by spatial transcriptomics. Science 353, 78–82 (2016).

6. Vickovic, S. et al. High-definition spatial transcriptomics for in situ tissue profiling. Nat. Methods 16, 987–990 (2019).

7. Rodriques, S. G. et al. Slide-seq: A scalable technology for measuring genome-wide expression at high spatial resolution. Science 363, 1463–1467 (2019).

8. Merritt, C. R. et al. Multiplex digital spatial profiling of proteins and RNA in fixed tissue. Nat. Biotechnol. 38, 586–599 (2020).

9. Butler, D. et al. Shotgun transcriptome, spatial omics, and isothermal profiling of SARS-CoV-2 infection reveals unique host responses, viral diversification, and drug interactions. Nat. Commun. 12, 1660 (2021).

10. Brady, L. et al. Inter- and intra-tumor heterogeneity of metastatic prostate cancer determined by digital spatial gene expression profiling. Nat. Commun. 12, 1426 (2021).

11. Desai, N. et al. Temporal and spatial heterogeneity of host response to SARS-CoV-2 pulmonary infection. Nat. Commun. 11, 6319 (2020).

12. Lui, J. H., Hansen, D. V. & Kriegstein, A. R. Development and Evolution of the Human Neocortex. Cel l vol. 146 332 (2011).

13. Kostovic, I. & Rakic, P. Developmental history of the transient subplate zone in the visual and somatosensory cortex of the macaque monkey and human brain. J. Comp. Neurol. 297, 441–470 (1990).

14. Zecevic, N., Chen, Y. & Filipovic, R. Contributions of cortical subventricular zone to the development of the human cerebral cortex. J. Comp. Neurol. 491, 109–122 (2005).

15. Nowakowski, T. J. et al. Spatiotemporal gene expression trajectories reveal developmental hierarchies of the human cortex. Science 358, 1318–1323 (2017).

16. Polioudakis, D. et al. A Single-Cell Transcriptomic Atlas of Human Neocortical Development during Mid-gestation. Neuron 103, 785–801.e8 (2019).

17. Pollen, A. A. et al. Molecular identity of human outer radial glia during cortical development. Cell 163, 55–67 (2015).

18. Huang, W. et al. Origins and Proliferative States of Human Oligodendrocyte Precursor Cells. Cell 182, 594–608.e11 (2020).

19. Monier, A. et al. Entry and distribution of microglial cells in human embryonic and fetal cerebral cortex. J. Neuropathol. Exp. Neurol. 66, 372–382 (2007).

20. Svensson, V., Teichmann, S. A. & Stegle, O. SpatialDE: identification of spatially variable genes. Nat. Methods 15, 343–346 (2018).

21. Willsey, A. J. et al. Coexpression networks implicate human midfetal deep cortical projection neurons in the pathogenesis of autism. Cell 155, 997–1007 (2013).

22. Satterstrom, F. K. et al. Large-Scale Exome Sequencing Study Implicates Both Developmental and Functional Changes in the Neurobiology of Autism. Cell 180, 568–584.e23 (2020).

23. Grasby, K. L. et al. The genetic architecture of the human cerebral cortex. Science 367, (2020).

24. Savage, J. E. et al. Genome-wide association meta-analysis in 269,867 individuals identifies new genetic and functional links to intelligence. Nat. Genet. 50, 912–919 (2018).

25. Thormann, A. et al. Flexible and scalable diagnostic filtering of genomic variants using G2P with Ensembl VEP. Nat. Commun. 10, 2373 (2019).

26. Andersson, A. et al. Single-cell and spatial transcriptomics enables probabilistic inference of cell type topography. Commun Biol 3, 565 (2020).

27. Kleshchevnikov, V., Shmatko, A., Dann, E. & Aivazidis, A. Comprehensive mapping of tissue cell architecture via integrated single cell and spatial transcriptomics. bioRxiv (2020).

28. Cable, D. M. et al. Robust decomposition of cell type mixtures in spatial transcriptomics. Nat. Biotechnol. (2021) doi: 10.1038/s41587-021-00830-w.

29. Hansen, D. V., Lui, J. H., Parker, P. R. L. & Kriegstein, A. R. Neurogenic radial glia in the outer subventricular zone of human neocortex. Nature 464, 554–561 (2010).

30. Kwan, K. Y., Sestan, N. & Anton, E. S. Transcriptional co-regulation of neuronal migration and laminar identity in the neocortex. Developmen t 139, 1535–1546 (2012).

31. Efremova, M., Vento-Tormo, M., Teichmann, S. A. & Vento-Tormo, R. CellPhoneDB: inferring cell-cell communication from combined expression of multi-subunit ligand-receptor complexes. Nat. Protoc. 15, 1484–1506 (2020).

32. Bhaduri, A. et al. Cell stress in cortical organoids impairs molecular subtype specification. Nature 578, 142–148 (2020).

33. Berg, S. et al. ilastik: interactive machine learning for (bio)image analysis. Nat. Methods 16, 1226–1232 (2019).

## References

[1] Pierre Boyeau, Romain Lopez, Jeffrey Regier, Adam Gayoso, Michael I Jordan, and Nir Yosef. Deep generative models for detecting differential expression in single cells. bioRxiv, page 794289, 2019.

[2] Shiqi Cui, Subharup Guha, Marco AR Ferreira, Allison N Tegge, et al. hmmseq: A hidden markov model for detecting differentially expressed genes from rna-seq data. The Annals of Applied Statistics, 9(2):901–925, 2015.

[3] Vitalii Kleshchevnikov, Artem Shmatko, Emma Dann, Alexander Aivazidis, Hamish W King, Tong Li, Artem Lomakin, Veronika Kedlian, Mika Sarkin Jain, Jun Sung Park, et al. Comprehensive mapping of tissue cell architecture via integrated single cell and spatial transcriptomics. bioRxiv, 2020.

[4] Alp Kucukelbir, Dustin Tran, Rajesh Ranganath, Andrew Gelman, and David M Blei. Automatic differentiation variational inference. The Journal of Machine Learning Research, 18(1):430–474, 2017.

[5] Hanna Mendes Levitin, Jinzhou Yuan, Yim Ling Cheng, Francisco JR Ruiz, Erin C Bush, Jeffrey N Bruce, Peter Canoll, Antonio Iavarone, Anna Lasorella, David M Blei, et al. De novo gene signature identification from single-cell rna-seq with hierarchical poisson factorization. Molecular systems biology, 15(2):e8557, 2019.

[6] Romain Lopez, Jeffrey Regier, Michael B Cole, Michael I Jordan, and Nir Yosef. Deep generative modeling for single-cell transcriptomics. Nature methods, 15(12):1053–1058, 2018.

[7] Davide Risso, Fanny Perraudeau, Svetlana Gribkova, Sandrine Dudoit, and Jean-Philippe Vert. A general and flexible method for signal extraction from single-cell rna-seq data. Nature communications, 9(1):1–17, 2018.

[8] John Salvatier, Thomas V Wiecki, and Christopher Fonnesbeck. Probabilistic programming in python using pymc3. PeerJ Computer Science, 2:e55, 2016.

[9] Abhishek K Sarkar and Matthew Stephens. Separating measurement and expression models clarifies confusion in single cell rna-seq analysis. BioRxiv, 2020.

[10] Oliver Stegle, Leopold Parts, Matias Piipari, John Winn, and Richard Durbin. Using probabilistic estimation of expression residuals (peer) to obtain increased power and interpretability of gene expression analyses. Nature protocols, 7(3):500, 2012.

[11] Shiquan Sun, Yabo Chen, Yang Liu, and Xuequn Shang. A fast and efficient count-based matrix factorization method for detecting cell types from single-cell rnaseq data. BMC systems biology, 13(2):1–8, 2019.

[12] F William Townes, Stephanie C Hicks, Martin J Aryee, and Rafael A Irizarry. Feature selection and dimension reduction for single-cell rna-seq based on a multinomial model. Genome biology, 20(1):1–16, 2019.

